# Assessing the utility of death assemblages as reference conditions in a common benthic index (M-AMBI) with simulations

**DOI:** 10.1101/2022.08.30.505344

**Authors:** Jansen A. Smith, Matthew J. Pruden, John C. Handley, Stephen R. Durham, Gregory P. Dietl

## Abstract

Incorporating paleontological data into the methods and formats already familiar to conservation practitioners may facilitate greater use of paleontological data in conservation practice. Benthic indices (e.g., Multivariate - AZTI Marine Biotic Index; M-AMBI) already incorporate reference conditions and are a good candidate for integration. In simulations of living communities under constant and changing environmental conditions, we evaluate the capacity of death assemblage reference conditions to replicate M-AMBI values when used in place of reference conditions from the final ten generations of the simulation or all five hundred simulated generations.

Reference conditions from all death assemblage scenarios successfully replicated correct remediation decisions in the majority of simulation runs with environmental change and stability. Variations in M-AMBI values were due to overestimated richness and diversity in the death assemblages but effects of changes to these parameters varied across scenarios, emphasizing the importance of evaluating multiple metrics. Time averaging was largely beneficial, particularly when environmental change occurred and short-term ecological observations (ten generations) produced incorrect remediation decisions. When the duration of time averaging is known, death assemblages can provide valuable long-term perspectives with the potential to outperform temporally constrained baseline information from monitoring the living community.

**Supplementary material:** All R code used to produce the simulation, analyze outputs, and create figures is available at: https://doi.org/10.5281/zenodo.6355921. The simulated data is also available at this location. Supplementary figures and analyses referred to in the text are available at the end of this document.

## INTRODUCTION

Paleontologists have begun to argue for the relevance of paleontological data in achieving conservation and restoration goals (Dietl and Flessa 2011, 2017; Louys et al. 2012; Kidwell 2013; Rick and Lockwood 2013; Dietl et al. 2015; Barnosky et al. 2017; Tyler and Schneider 2018). Nonetheless, paleontological data are rarely used in conservation practice, as conservation paleobiology struggles to navigate the spaces between research and implementation (Barnosky et al. 2017; Dietl et al. 2019; Smith et al. 2020). One strategy to increase the utility of conservation paleobiology data is to adapt it to the methods and data formats already familiar to decision makers (Dietl et al. 2016; Barnosky et al. 2017; Smith et al. 2020). Aquatic benthic indices, which describe the ecological quality of a water body based on the composition of the benthic (bottom dwelling) macroinvertebrate communities, present one such opportunity for conservation paleobiologists to use a metric already familiar to resource managers, due to the abundance of taxonomically identifiable remains of benthic organisms that lived at a site in the past, preserved as buried death assemblages on the sea floor (Nerlović et al. 2011; Leshno et al. 2015; Dietl et al. 2016; Leshno et al. 2016; Smith et al. 2020; Pruden et al. 2021). Here we use a simulated benthic community to investigate the differences in performance between death assemblage data and living assemblage data as reference condition information for the widely used marine benthic index, Multivariate AZTI Marine Biotic Index (M-AMBI).

### Benthic Indices and the Development of M-AMBI

In the 1940’s and 1950’s, environmental policy makers, managers, and decision makers (hereafter decision makers) recognized a need for easy and rapid assessments of water quality and pollution to enact effective water management legislation (Parran 1947; Weiss 1951; Hays 1987; Colten 2005). Initially, assessment methods were focused on measuring water chemistry and the impact of pollution on economic utility (LeBosquet 1950; Doudoroff and Warren 1957; Seabloom 1958; Smallhorst 1960). The importance of biotic dimensions, such as maintaining diverse ecological communities, was later recognized and incorporated in the valuation of water bodies (Gaufin and Tarzwell 1952, 1956; Woodiwiss 1964; Pratt and Coler 1976; De Pauw and Vanhooren 1983). Early water chemistry assessments were useful for determining levels of pollutants but not adequate to provide information on the response of the biota to environmental stressors (Metcalfe 1989), necessitating the development of new assessment methods.

Over several decades, researchers worked to develop biotic indices by categorizing marine macrobenthic species into groups based on their responses to environmental stressors, notably organic enrichment (see supplement for more details on the importance of organic enrichment in the history of biotic indices; Pearson and Rosenberg 1978; Gray 1979). One of the more common categorization schemes was proposed by Hily (1984), Gelmarec (1986) and Grall and Glemarec (1997) who divided macroinvertebrate species into five ecological groups (EG) according to increasing tolerance to organic enrichment: EGI (species sensitive to organic enrichment); EGII (species indifferent to enrichment and always present in low densities); EGIII (species tolerant to enrichment and whose populations are stimulated by organic enrichment); EGIV and EGV (second- and first-order opportunistic species, respectively, with short life cycles and adaptations to life in reduced sediments). In 2000, Borja et al. applied these ecological groupings in the AZTI Marine Biotic Index (AMBI) to calculate a score for the community as weighted proportions of the five EGs:

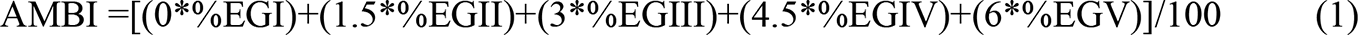

The resulting AMBI scores range from zero to seven and correspond to one of five Ecological Quality Statuses (EcoQS) – (‘High’, 0 < AMBI < 1.2; ‘Good’, 1.2 < AMBI < 3.3; ‘Moderate’, 3.3 < AMBI < 4.3; ‘Poor’, 4.3 < AMBI < 5.5; ‘Bad’, 5.5 < AMBI < 7; Borja et al. 2004) – which determines whether remediation is needed. For many applications, emphasis is placed on the Good-Moderate boundary, with sites classified as High or Good not requiring remediation, and those classified as Moderate, Poor, or Bad requiring remediation action (Vincent et al. 2002; Borja et al. 2004).

In 2000, a few months after the publication of AMBI, the European Commission published the European Water Framework Directive (WFD), which aimed to establish common principles and definitions of water quality (European Commission 2000). The WFD required European countries to adopt ecological quality indices incorporating indicator species, species richness, diversity, and abundance. The timing of the publication of AMBI and the WFD resulted in AMBI being adopted in ecological quality assessments throughout Europe (Borja et al. 2019). AMBI has subsequently been applied around the globe, with the original publication having been cited in articles from 84 countries as of 2022 (SCOPUS, March 9, 2022; Borja et al. 2019).

Despite its success, AMBI did not meet all requirements established by the WFD (Borja et al. 2019), so seven years later it was modified to incorporate species richness and Shannon-Wiener diversity via a factor analysis, creating the Multivariate-AMBI (M-AMBI; Muxika et al. 2007). The index scores from the factor analysis are placed along an orthogonal gradient from a user-defined reference condition to a highly degraded condition, and the resulting position within Euclidean space is the M-AMBI value. The M-AMBI values range from zero to one and correspond to one of the five EcoQS classifications (‘High’, 0.77 < M-AMBI < 1; ‘Good’, 0.53 < M-AMBI < 0.77; ‘Moderate’, 0.39 < M-AMBI < 0.53; ‘Poor’, 0.20 < M-AMBI < 0.39; ‘Bad’, 0 < M-AMBI < 0.20; Muxika et al. 2007). M-AMBI has been used globally, due largely to its incorporation of reference conditions, and it has been officially adopted for assessing ecological quality under the WFD (European Commission 2018; Reizopolou et al. 2018) by seven European countries (Bulgaria, France, Germany, Greece, Italy, Romania, Spain).

### Reference Conditions

Over the past several decades, government agencies have enacted legislation to establish reference conditions for water quality management (e.g., Canada-British Columbia Water Quality Monitoring Agreement; European Water Framework Directive). Reference conditions can be used to set restoration targets based on the past state of an ecosystem, to set future environmental targets to guide remediation actions, or as a benchmark for tracking changes in ecological quality (Gibson and Bowman 2000; Stoddard et al. 2006; Gann et al. 2019). Borja et al. (2012) recommended three methods for setting reference conditions: modeling expected biological attributes based on environmental variables, conditions from pristine or least-disturbed areas within the same ecoregion, and historical information such as prior assessments or model hindcasting.

Historical information is underutilized for setting reference conditions despite the growing recognition of the need for long-term (decadal to centennial scale) ecological data to assess changes in ecological quality from pollution, and to distinguish between natural and anthropogenic stressors (Wolfe et al. 1987; Krebs et al. 2001; Latimer et al. 2003; Lindenmayer et al. 2012). In contrast, long-term ecological data are used disproportionately more frequently for informing policy decisions than studies with shorter durations (less than a decade; Hughes et al. 2017). The significance of long-term data in both scientific assessments and policy decisions notwithstanding, a recent survey of authors from the US National Research Council highlighted the mismatch between the demand and availability for long-term data (Hughes et al. 2017). There are few long-term ecological studies that cover the length of time needed to encapsulate natural variability on decadal, centennial, and longer timescales (Lindenmayer et al. 2012; Dornelas et al. 2018; Estes et al. 2018).

Identifying appropriate historical reference conditions using existing assessments and monitoring records for benthic indices is challenging, particularly as no truly pristine sites likely exist (e.g., Halpern et al. 2008). The importance, yet lack of, historical reference conditions provides an opportunity for the incorporation of geohistorical data (Dietl et al. 2016; Smith et al. 2020; Pruden et al. 2021). In particular, molluscan death assemblages can provide location-specific, long-term ecological information on the response of marine benthic communities to anthropogenic stressors, which would otherwise be inaccessible to decision makers (Nerlović et al. 2011; Leshno et al. 2015; Dietl et al. 2016; Leshno et al. 2016; Smith et al. 2020; Pruden et al. 2021). Moreover, studies with data from Europe (Dietl et al. 2016) and the United States (Pruden et al. 2021) have demonstrated that mollusks are reliable proxies for the whole benthic macroinvertebrate communities under the AMBI and M-AMBI framework. Before the implementation of this approach with molluscan death assemblages, however, it is necessary to evaluate the effects of common death assemblages biases (e.g., preservation bias, time averaging) on the parameters used to define reference conditions (i.e., richness, diversity, AMBI), and their combined effects on M-AMBI (Dietl et al. 2016; Smith et al. 2020; Pruden et al. 2021).

### Death Assemblages

Death assemblages contain individuals that lived during the previous years to millennia (Kowalewski et al. 1998; Kidwell 2002; Kidwell and Tomasovych 2013) and because they often accumulate *in situ*, they also provide a location-specific record of past iterations of the community in the habitat (e.g., Tyler and Kowalewski 2017; Arkle and Miller 2018; Casebolt and Kowalewski 2018; Hyman et al. 2019). Through their spatial fidelity and extended temporal scope, death assemblages have the potential to provide a useful perspective for conservation and restoration decision making (e.g., Kidwell 2007; Albano and Sabelli 2011; Dietl and Durham 2016; Smith and Dietl 2016; Albano et al. 2017; Dietl and Smith 2017; Tyler and Kowalewski 2017; Wingard 2017; Hyman et al. 2019; Lockwood and Mann 2019; Powell et al. 2020). Before death assemblages and other sources of geohistorical data can fulfill this potential, it is critical to evaluate the consequences of taphonomic processes that may bias the accumulation of individuals. Though there is a robust literature on these processes and biases (e.g., Cummins et al. 1986; Davies et al. 1989; Kowalewski et al. 1998; Behrensmeyer et al. 2005; Lockwood and Chastant 2006; Olszewski and Kidwell 2007; Kosnik et al. 2009; Powell et al. 2011; Kidwell and Tomašovỳch 2013; Tomašovỳch et al. 2016; Smith et al. 2021), they have not been considered with respect to the use of benthic indices like M-AMBI (Smith et al. 2020).

Smith et al. (2020) identified six potential biases that need to be evaluated to improve the confidence associated with M-AMBI values calculated when using death assemblages as reference conditions. These biases include, (i) molluscan sensitivity to disturbance, (ii) proportion of living community that is mollusks, (iii) preservation of species abundance, (iv) increased richness in death assemblages, (v) increased evenness in death assemblages, and (iv) life span bias (Smith et al. 2020). The first two biases relate to characteristics of living communities, in which mollusks often represent only a fraction of the whole benthic community and have a tendency to be more sensitive to disturbance than other taxa. The next four biases are interconnected and related to taphonomic processes. Absolute abundances of taxa in death assemblages are often skewed with respect to abundances in living assemblages, although rank-order abundance in the death assemblage tends to be more resistant to taphonomic bias than absolute abundance when the environment is stable (e.g., Kidwell 2002, 2007). Many factors can influence abundances in the death assemblage (e.g., Cummins et al. 1986; Kidwell 2002, 2007; Olszewski and Kidwell 2007; Kosnik et al. 2009; Kidwell and Rothfus 2010; Powell et al. 2011; Olszewski 2012; Tomašovỳch et al. 2016, 2018; Smith et al. 2019, 2021), including differences in the life spans of species and the consequent rate at which dead individuals enter the death assemblage (Cronin et al. 2018). Thus, the relationship between preservation of absolute abundance and EcoQS is likely to be variable and dependent on characteristics of particular species in an assemblage. We refrain from incorporating life span bias here, as more data on molluscan life spans and the effects on death assemblage accumulation are needed (Kidwell and Rothfus 2010; Cronin et al. 2018), particularly with respect to M-AMBI.

The remaining two biases have been consistently reported as features of death assemblages due to time averaging: increases in richness and evenness relative to the corresponding living assemblage (Kidwell 2002, 2009, 2013; Olszewski and Kidwell 2007; Burkli and Wilson 2017). As the focus here is on M-AMBI, we focus on increased richness and on diversity, rather than evenness. Although Smith et al. (2020) did not discuss the potential effects of diversity on M-AMBI, we expect that the direction of the trend will be similar to that of evenness and richness. Indeed, the ratio of death assemblage to living assemblage diversity has been reported to be as high as 1.8 (Tomašovỳch and Kidwell 2010*a*). Likewise, with standardized sample size and no change in environmental conditions, the ratio of death assemblage to living assemblage species richness can range from 1.22 – 1.8 (Kidwell 2002 2009; Tomašovỳch and Kidwell 2010*a*). These values depend on the scale of sampling (habitat scale to point scale, respectively) and the degree of time averaging, with increases driven by the accumulation of rare species over the extended duration of sampling allowed by time averaging (Kidwell 2002, 2009; Tomašovỳch and Kidwell 2010*a*). Based on these well-documented taphonomic biases, we expect that using reference conditions from a death assemblage to calculate M-AMBI values for the corresponding living community will result in lower M-AMBI values than when using reference conditions drawn from preceding generations of the living community.

## METHODS

### Simulating death assemblages to quantify biases

We used a simulation approach to investigate the effects of time averaging on M-AMBI values when death assemblages were used to generate reference conditions instead of the living assemblage. The simulation used a version of the forward function from the ecolottery R package (Munoz et al. 2018); R code: https://doi.org/10.5281/zenodo.6355921, which was developed to perform forward-in-time simulations of community composition that incorporate life history variables and environmental filters. The function allowed us to retain a complete census of a living community over time and was modified to also retain information on dead individuals, which facilitated comparisons between the living community and the resulting death assemblage. The model focused on the flow of mollusk individuals over time between three simulation components: metacommunity, local community, and death assemblage. The metacommunity was composed of 100,000 individuals distributed uniformly across 100 species (i.e., 1,000 individuals of each species). The local community was a random subset of the metacommunity, and the death assemblage was composed of the local community individuals that died over the course of the simulation. At the start of each simulation run (n = 100 runs for each scenario), each species was assigned values for three characteristics: ecological group, ecological preference, and preservation potential. The values were assigned separately for each simulation run to allow them to vary at random with respect to each other. Together, these three characteristics interacted with terms for immigration probability and an environmental filter to determine the species composition of the individuals flowing between the three model components over the course of the simulation. To set up our comparison of M-AMBI values based on living community or death assemblage reference conditions, we simulated both a constant environment scenario with a constant environmental filter and an environmental change scenario in which the environmental filter shifted part-way through the simulation.

The ecological group of each species was assigned by taking a random draw from the distribution of ecological groups for mollusks in the AMBI package v6.0 (supplementary material Figure S1; http://ambi.azti.es; Species List v.Dec2020; Borja et al. 2012). Thus, we made the simplifying assumptions that the metacommunity species composition was constant over time and was a random sample of all possible mollusk species (given the species list).

Because a species’ ecological group categorization is based on its ecology, the value for each species’ ecological preference characteristic (assumed here to represent the entirety of a species’ niche) was assigned based on the ecological group of the species. Mean ecological preference values were defined for each species by drawing a random value from the distribution of tolerance values for taxa expected to be found in each ecological group according to the theoretical model (Hily 1984; Hily et al. 1986; Majeed 1987; Grall and Glémarec 1997) used in the original construction of AMBI (Borja et al. 2000; supplementary material Figure S2). To incorporate within species variation, individuals (n = 1,000) of each species were assigned a value with a 0.01 standard deviation around the mean for the species according to a truncated normal distribution (min = 0, max = 1). The ecological preference trait values ranged from 0 to 1, where low values indicated a species was more sensitive (e.g., ecological group I) and high values indicated tolerance and opportunism (e.g., ecological group V). Because the percentage of mollusks classified as ecological group V is exceedingly small (< 1%; supplementary material Figure S1), the highest ecological group used in the simulation was IV.

The preservation potential for each species was assigned by drawing a value from a truncated normal distribution (mean = 0.01, sd = 0.005, min = 0.001, max = 0.02; following Tomašovỳch et al. 2016). Similar to ecological preference, the preservation potential value for individuals of a species was allowed to vary around the species value, but with a standard deviation of 0.0001. Smaller values for the preservation potential indicated a greater likelihood of being removed from the death assemblage when preservational bias was included in the simulation. Assignment of the preservation potential was random with respect to ecological group and ecological preference.

The starting point for the local community in each simulation run was a random sample of 250 individuals from the metacommunity. Each subsequent generation was determined by the forward function (Munoz et al. 2018); see Supplement 2 for R code) and parameterization of the simulation. The simulation was run for 550 generations (i.e., years) and the first 50 were removed prior to analysis to account for stochasticity in the early generations (i.e., model burn-in). At each step, a death rate of 0.40 was applied to generate turnover – increased from 0.33 used by Tomašovỳch and Kidwell (2010*a*) to reflect higher reported rates in the literature (e.g., Levinton and Bambach 1970; Powell et al. 1984; Prince et al. 1988). Dead individuals (n = 100 for each generation) were replaced with reproduction from the local community or selection from the metacommunity according to the immigration parameter (m = 0.1; i.e., likelihood of immigration; (Tomašovỳch and Kidwell 2010*a*). In both cases, the new individual was selected based on its ecological preference value and the predefined environmental filter distribution. The closer the value of the ecological preference value to the mean of the environmental filter distribution, the more likely an individual was to be included in the next generation of the community. Under the scenario for environmental stability, the environmental filtering distribution was defined by mean = 0.20 and standard deviation = 0.05 with the intention of generating a community corresponding to relatively high EcoQS. The scenario for environmental change began with the same environmental filtering distribution, but the mean was shifted to 0.80 for the final 100 generations of the simulation to represent an abrupt degradation of the environment.

Over the course of each simulation run, every individual that died was passed to the death assemblage, which was sampled under four general scenarios after the simulation was completed. The four scenarios included: (1) perfect preservation (i.e., no preservational bias or effects of time averaging) where every dead individual had an equal probability of being sampled; (2) preservational bias where sampling probability depended on the preservation potential value assigned to each species; (3) time averaging bias where sampling probability was higher for more recently dead individuals; (4) preservational and time averaging bias where sampling probability depended on both the preservation potential and time since death for each individual (Table 1). For each scenario, ten random samples of 250 individuals each were taken from the death assemblage list in order to mimic the tendency for counts of individuals in death assemblages to be an order of magnitude larger than the living assemblage to which they are compared.

**Table 1.** Death assemblage scenarios, the features they include, and the names used for their reference.

|  | Preservational<br>Bias | Time Averaging | k-values | DARC Name |
| --- | --- | --- | --- | --- |
| DA Scenario 1 | No | Uniform | NA | Perfect |
| DA Scenario 2 | Yes | Uniform | NA | Preservation |
| DA Scenario 3 | No | Slightly Skewed Young | 0.5 | High TA |
|  | No | Moderately Skewed Young | 1.0 | Medium TA |
|  | No | Greatly Skewed Young | 1.5 | Low TA |
| DA Scenario 4 | Yes | Slightly Skewed Young | 0.5 | High TA w/ Pres |
|  | Yes | Moderately Skewed Young | 1.0 | Medium TA w/ Pres |
|  | Yes | Greatly Skewed Young | 1.5 | Low TA w/ Pres |

When simulating a death assemblage with non-uniform time averaging (i.e., DA scenarios 3 and 4), the likelihood of an individual being sampled was defined by a right-censored Weibull distribution (following equation 6 from Tomašovỳch et al. 2016):

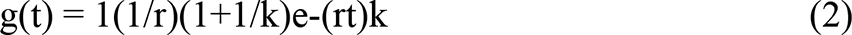

where *r* is the rate of decay and corresponds with our preservation potential values, *t* is the age of the individual (i.e., 500 minus the generation number in which it died), and *k* is a parameter determining whether the loss rate decreases (k < 1) or increases (k > 1) over time. If *k* = 1, the loss rate did not change over time (i.e., one-phase exponential decay, (Tomašovỳch et al. 2016). Because we were interested in understanding the effects of time averaging on M-AMBI results when using a death assemblage reference condition, we tested three values for k (0.5, 1.0, 1.5) in our sampling for DA scenarios 3 and 4. For DA scenario 3, which did not include preservational bias, the mean preservation potential value for all species (i.e., 0.01) was used for the *r* parameter.

### Calculating and comparing M-AMBI values

#### M-AMBI with different reference conditions

As the focal point of this study, we evaluated changes to M-AMBI values in the last generation of the simulation (i.e., the “living community”) when using different reference conditions. To establish reference conditions, richness, evenness, and AMBI were calculated for each generation in the simulations and for each of the death assemblage samples. For both environmental scenarios, ten different reference conditions were used: two live assemblage reference conditions (LARCs) and eight death assemblage reference conditions (DARCs). Reflecting perfect knowledge, the first LARC used the optimal values (i.e.., highest richness, highest diversity, lowest AMBI) recorded for each of the parameters at any point in the simulation (e.g., the highest richness could occur at generation 154 and the lowest AMBI at generation 379). The second LARC used optimal parameter values from only the final ten generations of the living community to more closely approximate an amount of information that a biological monitoring program might have. The eight DARCs were based on the death assemblages from the scenarios described above (Table 1). The optimal parameter values were taken from the ten death assemblage samples “collected” after each run of the simulation.

#### Fidelity of M-AMBI values

The M-AMBI values calculated when using the death assemblage scenarios as reference conditions were compared to those based on the two living community reference condition scenarios with respect to the Good-Moderate boundary because it is typically the most important threshold for decision making (see Supplement 1 for all values for all EcoQS). We evaluated the fidelity of the living-community- and death-assemblage-based results using recall, precision, and F1 (see also Pruden et al. 2021 Figure 1). Recall is the probability that our model returned the same remediation decision for the final living generation from the simulation when using as reference conditions either the ten generations LARC or a DARC in place of the perfect knowledge LARC for a given simulation (i.e., the probability that if the last generation required remediation according to the perfect LARC, the same decision was reached using one of the other reference conditions). Precision is the probability that the remediation decision for the last generation of the simulation when using the ten generation LARC or a DARC as the reference condition, is correct with regards to the perfect knowledge LARC (i.e., if the last generation was classified as requiring remediation using one of the other reference conditions, what is the probability that this decision was also reached using the perfect knowledge LARC scenario). F1 is the harmonic mean of recall and precision, and is calculated as 2*((Recall * Precision)/(Recall + Precision)) (Goutte and Gaussier 2005; Figure 1).

**Figure 1.**
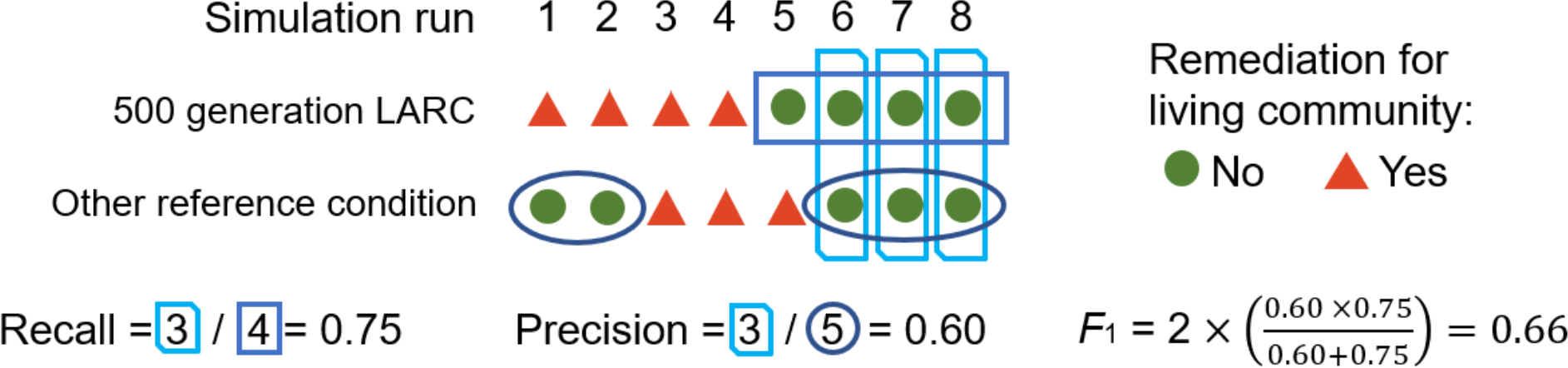
Schematic representation of the calculation of recall, precision, and F1 from hypothetical simulation runs. Adapted from Pruden et al. (2021).

#### Statistical analysis

To understand the effects of the preservation and time averaging biases on the calculation of M-AMBI values, we evaluated the seven death assemblage scenarios (DA Scenarios 2 - 4) that included variations in these parameters against the scenario with a perfect death assemblage (DA Scenario 1; Table 1). For both environmental scenarios, the differences between the M-AMBI values calculated using preceding generations of the living community (both five hundred and ten generations) as reference conditions and using the death assemblage as reference conditions were evaluated using a Kruskall-Wallis test with the Holm p-value adjustment method to control the familywise error rate (Aickin and Gensler 1996; Hecke 2012; see supplementary material for discussion of test choice).

Parameter values in DARCs used to calculate M-AMBI values exhibited characteristic biases (e.g., increased richness). and we evaluated the sensitivity of M-AMBI values to these biases. After quantifying differences in the parameter values between each of the two LARCs and eight DARCs, the differences were standardized using z-scores and fitted with a generalized linear mixed model. Regression coefficients from these models were interpreted as the change expected for differences in M-AMBI values due to a one standard deviation change in differences of parameter values between the reference conditions being compared. That is, the coefficients represent the sensitivity of M-AMBI values to changes in DARC parameter values relative to their LARC counterparts.

## RESULTS

### Constant Environment

In the set of simulation runs with a constant environment, the EcoQS categorizations of the final generation of the simulation (i.e., the “living community”) were unchanged regardless of the reference condition used to calculate the M-AMBI values (Figure 2). The three fidelity metrics used to evaluate the potential of the death assemblage to provide M-AMBI reference conditions showed complete agreement in EcoQS and, therefore, the implied remediation decision, with respect to the perfect LARC (i.e., reference conditions derived from all five hundred generations of the LA; see supplementary material Tables S1 and S2 for fidelity metric scores by EcoQS).

**Figure 2.**
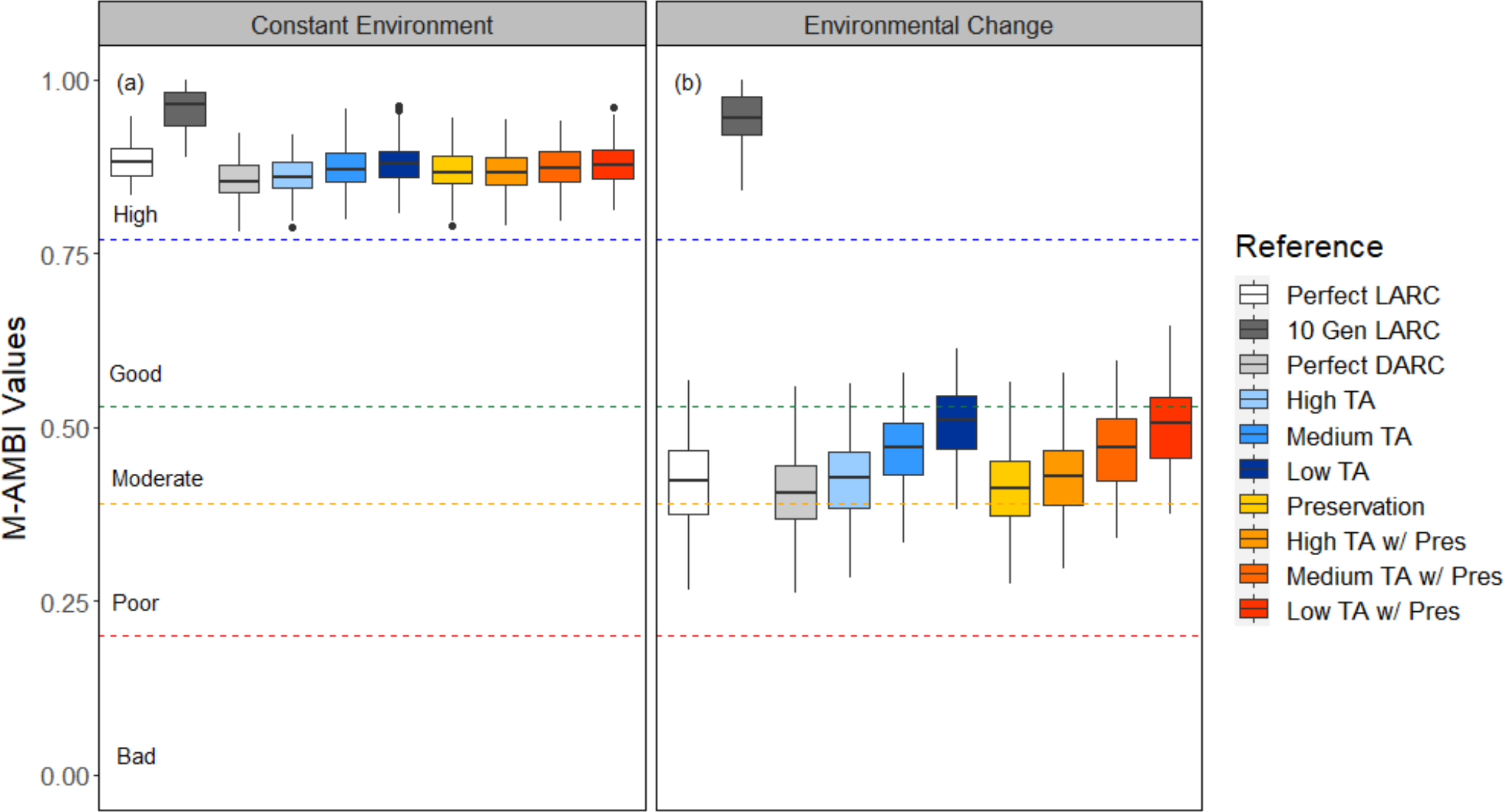
M-AMBI values for the final generation (i.e., the “living community”) of the constant (a) and changing (b) environment simulations, calculated using ten reference condition scenarios. n = 100 runs per environment simulation. Dashed lines delineate Ecological Quality Status boundaries: ‘High’, 0.77 < M-AMBI < 1; ‘Good’, 0.53 < M-AMBI < 0.77; ‘Moderate’, 0.39 < M-AMBI < 0.53; ‘Poor’, 0.20 < M-AMBI < 0.39; ‘Bad’, 0 < M-AMBI < 0.20; Muxika et al., 2007).

Compared to all DARC cases, the differences between M-AMBI values calculated using only the ten generation LARC were greater than when using the five hundred generation LARC (i.e., “perfect knowledge”; Figure 3). In either LARC case, M-AMBI values calculated with reference conditions drawn from the perfect death assemblage (DA Scenario 1) were most different (Figure 3; Table 3). Differences between M-AMBI values calculated with the perfect DARC and the LARCs were used as the null condition to compare the effects of preservation and time averaging when they were included to generate the other death assemblage reference conditions (DA Scenarios 2 - 4; Table 1). Compared with this null condition, all other DARCs produced significantly different differences with the exception of DA scenario 3 with k = 0.5 (Table 4). Making the same set of comparisons with the ten generation LARC, only DA scenarios 3 and 4 with k = 0.5 were not significantly different from the perfect DARC (Table 4). Increasing values of k (i.e., decreasing time averaging, supplemental material Table S3) always resulted in increasing differences in M-AMBI values relative to the M-AMBI result calculated using the perfect DARC. Correspondingly, M-AMBI values based on DARCs with lower time averaging resulted in M-AMBI values that were more similar to those calculated using the LARCs than those based on DARCs with higher time averaging (Figure 3).

**Figure 3.**
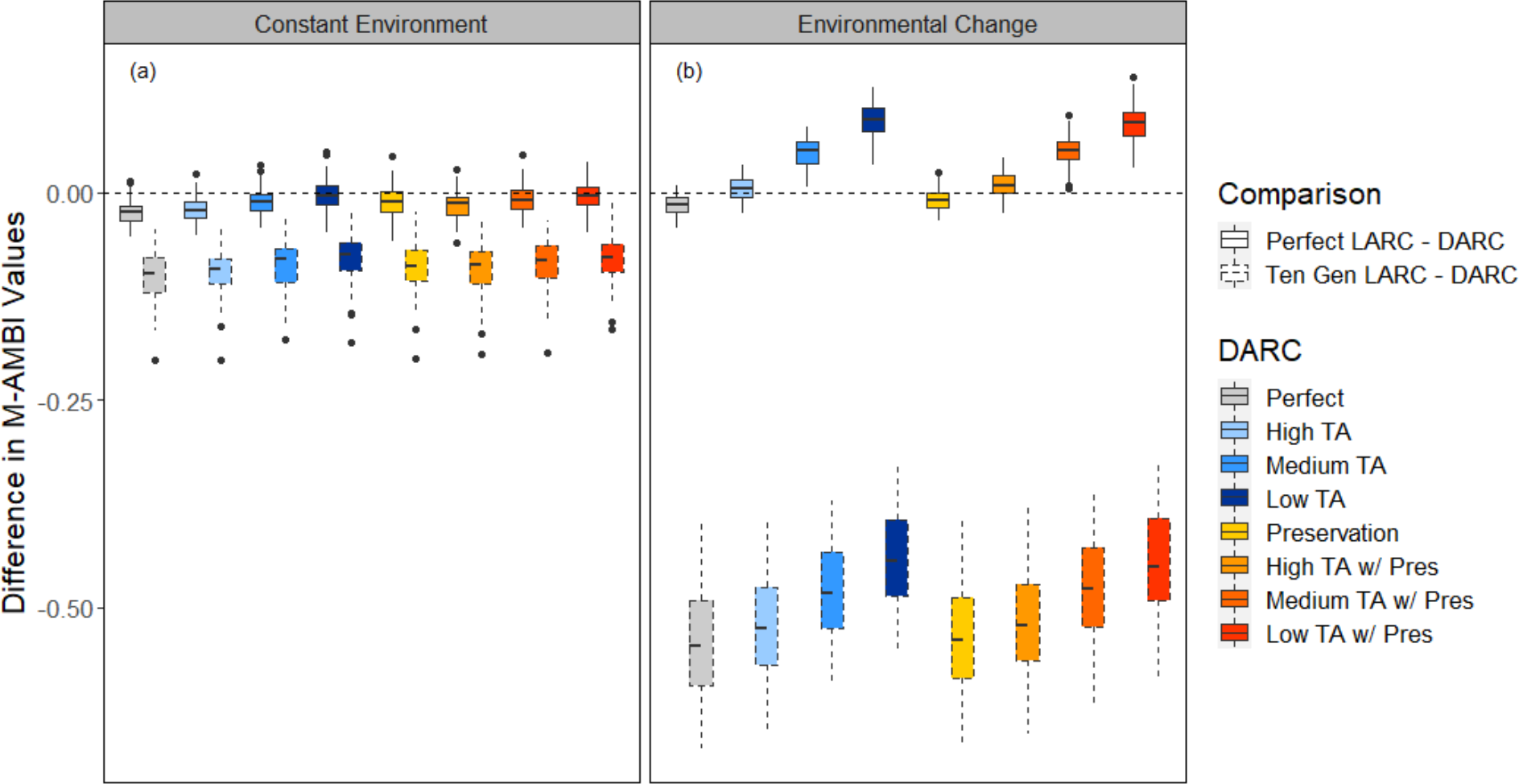
Differences in the calculated M-AMBI values of final generation of the simulation (i.e., the “living community”) for (a) a constant environment and (b) a changing environment when using reference conditions from the death assemblage rather than from all generations (Perfect LARC - DARC) and the last ten generations (Ten Gen LARC - DARC) of the living community. LARC = Live assemblage reference condition; DARC = Death assemblage reference condition; TA = time averaging; w/Pres indicates the inclusion of between species differences in preservation potential.

**Table 2.** Fidelity metric scores for capacity of M-AMBI results based on different reference conditions to identify the correct remediation decision* (relative to the M-AMBI result based on the perfect live assemblage reference condition). High metric values are bolded in the environmental change simulation columns.

| Reference | Remediation Recommended | Constant Environment |  |  | Environmental Change |  |  |
| --- | --- | --- | --- | --- | --- | --- | --- |
|  |  | Recall | Precision | F1 Score | Recall | Precision | F1 Score |
| 10 Generation LARC | No | 1.00 | 1.00 | 1.00 | **1.00** | 0.03 | 0.06 |
|  | Yes | NA | NA | NA | 0.00 | NA | NA |
| Perfect DARC | No | 1.00 | 1.00 | 1.00 | 0.33 | **1.00** | 0.50 |
|  | Yes | NA | NA | NA | **1.00** | **0.98** | **0.99** |
| High TA | No | 1.00 | 1.00 | 1.00 | **0.67** | **1.00** | **0.80** |
|  | Yes | NA | NA | NA | **1.00** | **0.99** | **0.99** |
| Medium TA | No | 1.00 | 1.00 | 1.00 | **1.00** | 0.50 | **0.67** |
|  | Yes | NA | NA | NA | **0.97** | **1.00** | **0.98** |
| Low TA | No | 1.00 | 1.00 | 1.00 | **1.00** | 0.08 | 0.15 |
|  | Yes | NA | NA | NA | **0.64** | **1.00** | **0.78** |
| Preservation | No | 1.00 | 1.00 | 1.00 | **0.67** | **1.00** | **0.80** |
|  | Yes | NA | NA | NA | **1.00** | **0.99** | **0.99** |
| High TA w/ Pres | No | 1.00 | 1.00 | 1.00 | **1.00** | **1.00** | **1.00** |
|  | Yes | NA | NA | NA | **1.00** | **1.00** | **1.00** |
| Medium w/ Pres | No | 1.00 | 1.00 | 1.00 | **1.00** | 0.27 | 0.43 |
|  | Yes | NA | NA | NA | **0.92** | **1.00** | **0.96** |
| Low TA w/ Pres | No | 1.00 | 1.00 | 1.00 | **1.00** | 0.08 | 0.15 |
|  | Yes | NA | NA | NA | **0.66** | **1.00** | **0.80** |

\*Metric values for all EcoQS categories are provided in supplementary material Tables S1 and S2.

**Table 3.**
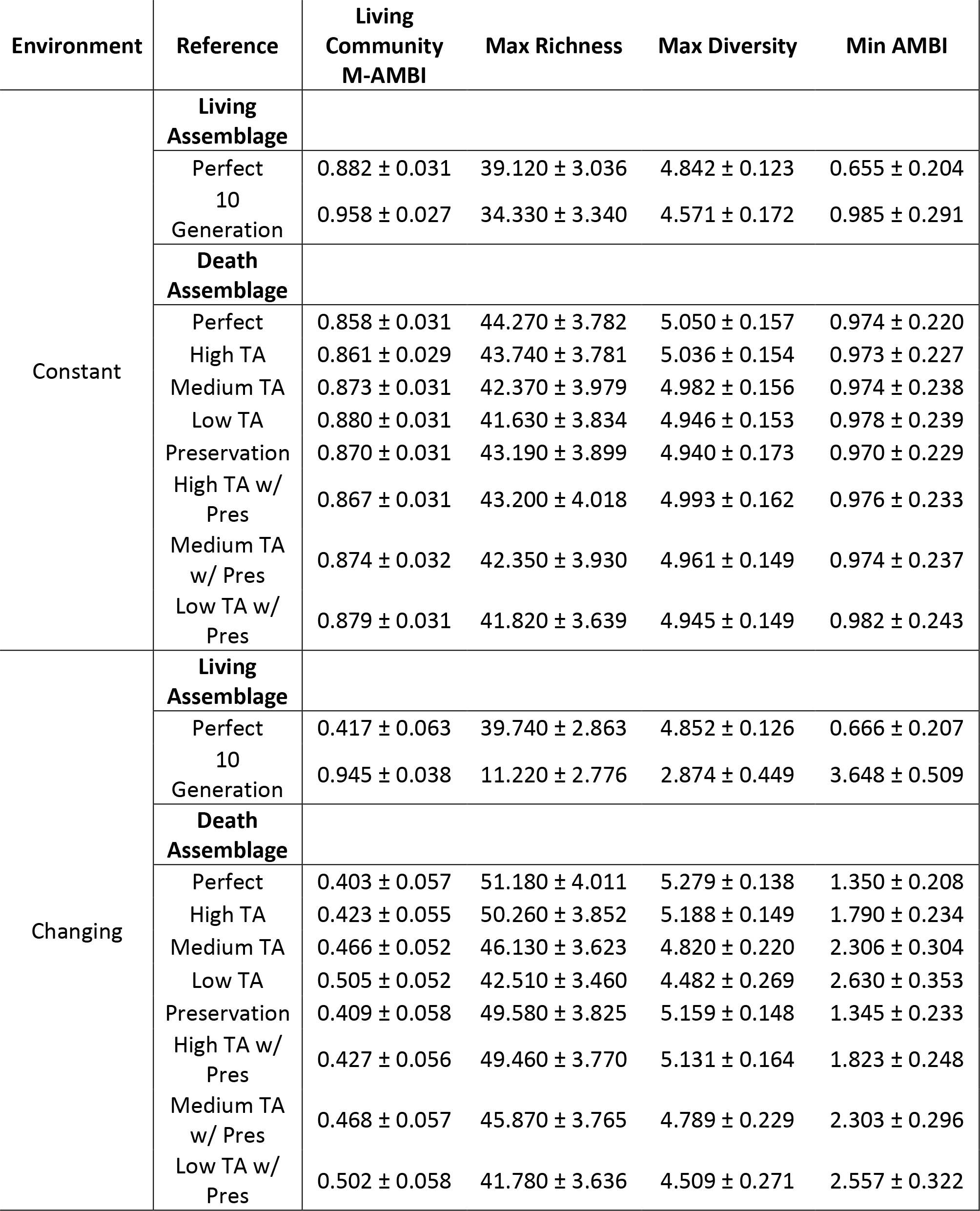
M-AMBI values of the living community (i.e., final simulation generation) and parameters used as reference conditions averaged across the one hundred simulation runs.

**Table 4.**
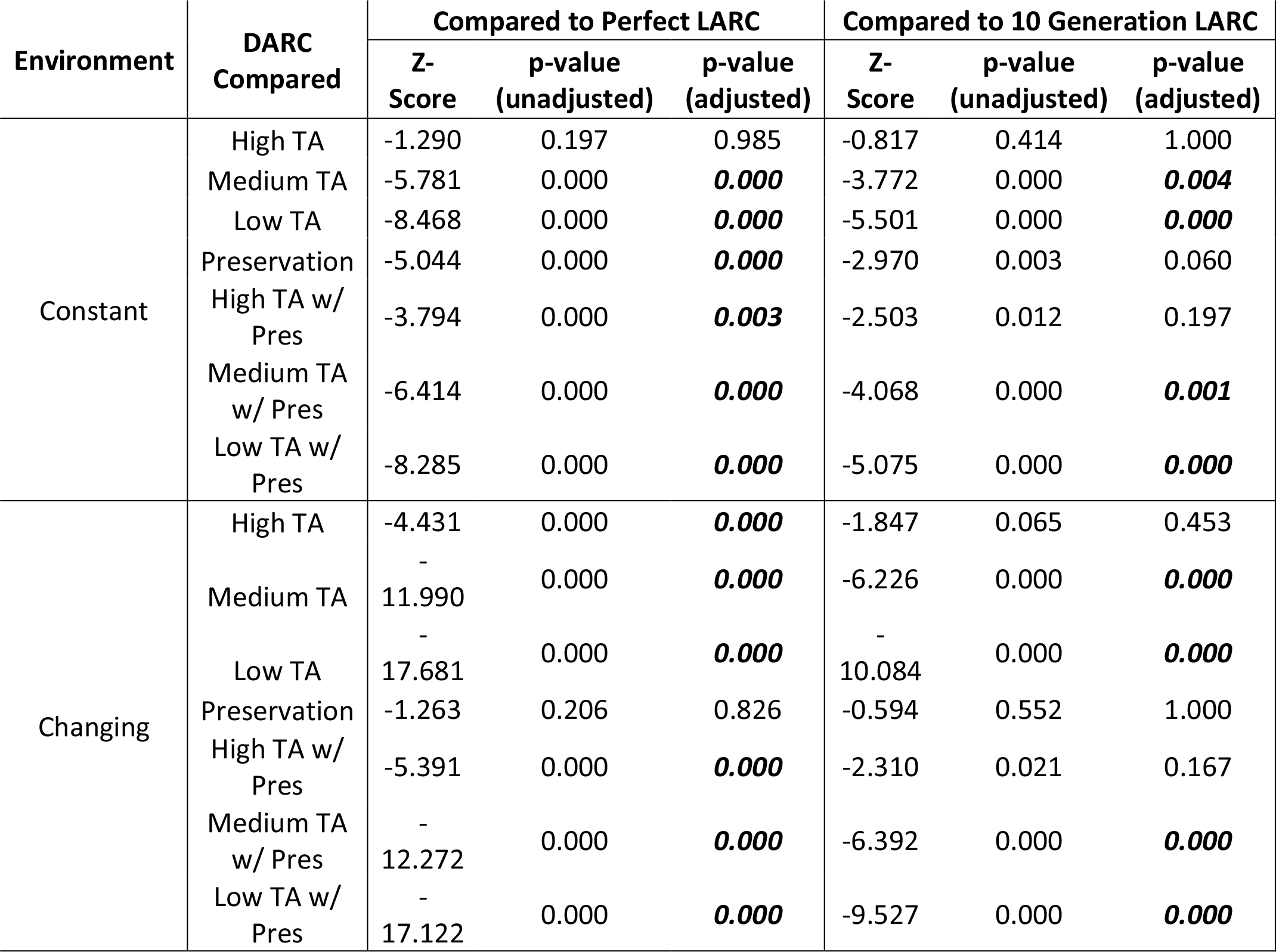
Results of the Kruskal-Wallis analysis. In both environmental scenarios, the difference between each death assemblage reference condition (DARC Compared) and live assemblage reference condition (LARC) was compared to the difference between the perfect DARC and the respective LARC. Significant (adjusted p-value < 0.05) differences are bolded.

Two of the three parameters used to establish reference conditions were sensitive to time averaging in the constant environment simulations. With and without preservation bias, increases in time averaging (i.e., decreasing k values) corresponded to increases in richness and diversity (Table 3). As the k-values decreased, richness and diversity values approached the values calculated for the perfect DARC. The magnitude of these changes was relatively small, as the greatest difference in mean values between scenarios (richness = 2.64, diversity = 0.104) was less than the standard deviation on the parameter means (Table 3). The magnitude of differences between DARC parameter values collectively and the individual LARC parameter values was larger. The perfect LARC parameter values, while still lower than those in the death assemblage, were higher than those calculated for the ten generation LARC. AMBI values calculated for the DARCs were relatively stable. The ten generation LARC AMBI value was the highest, but only marginally compared to the DARC values. The lowest AMBI value used in the reference conditions was from the perfect LARC.

Changes to the richness values used in the reference conditions, when switching from one of the LARCs to any of the DARCs, had the largest effect on the M-AMBI values (Table 5). The relative sensitivity of the M-AMBI values to diversity and AMBI values switched when comparing to the perfect and ten generation LARCs. AMBI had a larger effect when switching to a DARC from the perfect LARC whereas changes in diversity values had a greater impact when switching from the ten generation LARC (Table 5). In either case, the effect of changes to the parameter values in the DARCs was constant across the eight DARCs within comparisons to the individual LARCs.

**Table 5.** Sensitivity of M-AMBI values to changes in the three parameters used to establish reference conditions when switching from a living assemblage reference condition (LARC) to a death assemblage reference condition (DARC). Values have been normalized to be directly comparable in their effect size. Note: Sensitivity to parameters was identical within LARC comparisons when the environment was constant, so the values are only given once.

| Environment | LARC | DARC | Richness | Diversity | AMBI |
| --- | --- | --- | --- | --- | --- |
| Constant | Perfect | All | -0.013 ± 0.000 | -0.005 ± 0.000 | 0.007 ± 0.000 |
|  | 10<br>Generation | All | -0.018 ± 0.000 | -0.009 ± 0.000 | 0.012 ± 0.000 |
| Changing | Perfect | Perfect | -0.008 ± 0.001 | -0.022 ± 0.002 | 0.008 ± 0.002 |
|  |  | High TA | -0.008 ± 0.001 | -0.021 ± 0.002 | 0.006 ± 0.002 |
|  |  | Medium TA | -0.009 ± 0.001 | -0.016 ± 0.001 | 0.008 ± 0.001 |
|  |  | Low TA | -0.011 ± 0.001 | -0.012 ± 0.001 | 0.011 ± 0.001 |
|  |  | Preservation | -0.007 ± 0.001 | -0.020 ± 0.002 | 0.012 ± 0.002 |
|  |  | High TA w/<br>Pres | -0.009 ± 0.001 | -0.020 ± 0.002 | 0.007 ± 0.002 |
|  |  | Medium TA<br>w/ Pres | -0.009 ± 0.001 | -0.015 ± 0.001 | 0.010 ± 0.001 |
|  |  | Low TA w/<br>Pres | -0.011 ± 0.001 | -0.012 ± 0.001 | 0.011 ± 0.001 |
|  | 10<br>Generation | Perfect | -0.008 ± 0.004 | -0.034 ± 0.003 | 0.046 ± 0.003 |
|  |  | High TA | -0.007 ± 0.004 | -0.039 ± 0.003 | 0.052 ± 0.004 |
|  |  | Medium TA | -0.011 ± 0.004 | -0.046 ± 0.004 | 0.057 ± 0.005 |
|  |  | Low TA | -0.013 ± 0.004 | -0.050 ± 0.005 | 0.060 ± 0.006 |
|  |  | Preservation | -0.009 ± 0.004 | -0.035 ± 0.003 | 0.045 ± 0.003 |
|  |  | High TA w/<br>Pres | -0.008 ± 0.004 | -0.039 ± 0.003 | 0.050 ± 0.004 |
|  |  | Medium TA<br>w/ Pres | -0.010 ± 0.004 | -0.045 ± 0.004 | 0.055 ± 0.004 |
|  |  | Low TA w/<br>Pres | -0.011 ± 0.004 | -0.049 ± 0.005 | 0.059 ± 0.005 |

### Changing Environment

The EcoQS categorizations of the final generation of the simulation (i.e., the “living community”) were variable depending on the reference conditions used to calculate M-AMBI in the simulation runs with environmental change (Figure 2). Comparing to the five hundred generation LARC (i.e., perfect knowledge), the ten generation LARC showed the greatest difference (Figure 2; Table 3). The recall of the ten generation LARC was 0.00, indicating that using this reference condition resulted in the wrong remediation decision in every instance where the perfect LARC suggested that remediation was required (Table 2). Conversely, the “NA” value for remediation precision was because the ten generation LARC did not indicate a need for remediation in any simulation runs. By contrast, the DARC scenarios identified the “correct” decision for remediation in 64-100% of simulation runs (Table 2).

In the few simulation runs (n = 3) where the M-AMBI value for the last generation was categorized above the remediation boundary (i.e., “Good”) using the perfect LARC, the ten generation LARC did correctly recover those instances. The perfect DARC only recalled one of three such cases; however, the remaining DARCs recalled at least two (Table 2). The four DARCs with the most time averaging (i.e., uniform or k = 0.5, Table 1) consistently performed highest with all three fidelity metrics (Table 2). The high precision scores resulting from the use of these DARCs confirm the correct remediation decisions would be reached based on the categorizations produced from these DARCs, with respect to the perfect LARC. As evidenced by the perfect scores (i.e., 1.0) on each metric, the DA scenario with preservation and a longer duration of time averaging (k = 0.5) performed best overall.

As with the set of simulation runs under a constant environment, the differences in M-AMBI values calculated using DARCs were larger in comparison to the ten generation LARC than to the perfect LARC (Figure 3; Table 3). M-AMBI values calculated from the DARCs constructed with uniform time averaging (DA Scenarios 1 and 2) were statistically indistinguishable from each other (Table 4) and nearly indistinguishable from M-AMBI values calculated with the perfect knowledge LARC (Figure 3). Differences between the perfect DARC and remaining DARCs (DA Scenarios 3 and 4) in their differences with the perfect LARC were all statistically significant (Table 4). As compared to the ten generation LARC, the DARC scenarios with k = 0.5 were not significantly different from the perfect DARC but all others were (Table 4). With and without preservation bias and in comparisons to either LARC, increasing values of k (i.e., decreasing time averaging, supplemental Table S3) resulted in increasing distance from M-AMBI values calculated using the perfect DARC (Figure 2). As time averaging decreased, M-AMBI values calculated from DARCs were increasingly overestimated in comparison to the perfect LARC (Figure 2).

The parameter values used to define the reference conditions for M-AMBI calculations were all affected by the degree of time averaging in the death assemblage in the set of simulations with environmental change one hundred generations before the living community. Each of the parameters reached its most optimal value in the perfect DARC, which had uniform time averaging. As time averaging skewed the death assemblages towards the recent generations (i.e., k values increased) where the environmental filter was set to lower EcoQS conditions, richness and diversity decreased and AMBI increased. The inclusion of preservational bias had only a small effect on the parameters and ultimately the M-AMBI values. Of the ten sets of reference conditions used, the ten generation LARC was most different from the others: richness was four-fold lower, diversity values dropped by nearly half, and the minimum observed AMBI value was six times greater than in the perfect LARC (Table 3). The average M-AMBI value calculated based on the ten generation LARC was 0.945, however, categorizing the living community as High EcoQS and indicating no need for remediation (Figure 2).

When switching the reference conditions used to evaluate M-AMBI from one of the LARCs to one of the DARCs, there was a large degree of variability in how changes to the reference condition parameters affected M-AMBI values (Table 5). When switching from the perfect LARC to a DARC, changes to diversity values had the largest effect; however, the sensitivity of live community M-AMBI values to changes in the parameters became effectively identical as the extent of time averaging was reduced. The sensitivity to changes in diversity lessened whereas sensitivity to richness and AMBI increased. When switching from the ten generation LARC to a DARC, changes to the AMBI value used in the reference condition had the largest effect on M-AMBI values, followed by diversity. The sensitivity to richness was more than four times lower than to AMBI and diversity, comparing within each DARC. As the duration of time averaging decreased, the sensitivity of the M-AMBI values to changes in each of the parameters increased (Table 5).

## DISCUSSION

The outputs of the simulation framework developed here suggest that death assemblages can be appropriate for use as reference conditions in calculations of M-AMBI. Under scenarios with and without environmental change, death assemblage reference conditions (DARCs) faithfully reproduced the remediation decisions made using a perfect knowledge LARC when being used in their place to calculate M-AMBI values (i.e., last generation of the simulation). Moreover, DARCs consistently provided better estimates of correct M-AMBI values (i.e., those calculated with the perfect knowledge LARC) than the ten generation LARC (Figures 2 and 3), which was used as a more realistic approximation of the temporal duration of modern monitoring efforts. The effect of time averaging on M-AMBI values was minimal when the environment was constant; however, in the more realistic scenario where the environment changed, the time averaging in the death assemblage preserved the pre-environmental-change signal (Table 3) and was a factor in reproducing correct remediation decisions (Table 2). Preservation potential, which was assigned at random in this simulation framework, did not strongly influence the M-AMBI results, but did have minor effects on the parameters used to calculate reference conditions from the death assemblages. Though the simulation framework used here has some constraints, the results suggest that death assemblages can be a valuable tool with an unique temporal perspective for making remediation decisions in modern marine ecosystems.

### Death assemblages as reference conditions

Using reference conditions from time averaged death assemblages to calculate M-AMBI values returned results that would indicate the same remediation decision as the perfect LARC (hereafter, “correct remediation decision”) in all cases when the environment was constant and in the majority of cases (e.g., values ⩾ 64% for Recall = 15/16, Precision = 12/16, F1 = 12/16) when the environment changed (Table 2). We discuss these DARC results in order of increasing complexity used to generate the death assemblages (Table 1); considering the perfect DARC then the additions of preservational bias and time averaging.

In the simulation framework, the perfect death assemblage scenario allowed for an equal likelihood that any individual that died during the simulation run could be included in the death assemblage samples (perfect DARC). Like the perfect LARC, this perfect DARC was not realistic but provided a benchmark for evaluating the effects of preservational bias and time averaging in the other DA scenarios (Table 1). As expected (e.g., Kidwell 2002, 2009; Olszewski and Kidwell 2007; Tomašovỳch and Kidwell 2010; Tomašových and Kidwell 2011; Bürkli and Wilson 2017), the perfect DARC had higher richness and diversity than any other reference condition in both environmental scenarios (Table 3). Thus, the M-AMBI values based on the perfect DARC were the most underestimated in comparison to those that used the perfect LARC, regardless of environmental scenario (Figures 2 and 3).

The effect of uniform time averaging in the perfect DARC on AMBI values was directly related to the environmental scenario. As AMBI values are a weighted average of the ecological groups of the individuals in a sample (Equation 1; Borja et al. 2000), inclusion of an environmental change to simulate degradation resulted in less optimal AMBI values than when the environment was constant (Table 3). With a constant environment, time averaging was less important because individuals of all ages came from communities composed of species belonging to lower ecological groups and the consistently low AMBI values in all DARCs reflect this (Table 3). When environmental change was introduced one hundred generations before the end of the simulation, the composition of the living communities shifted gradually to include more species from higher ecological groups (supplementary Figure S7 - S9). The ecological signal in the perfect death assemblage lagged behind these changes to the living community, however, as the addition of new dead individuals was diluted by the abundance of individuals from low ecological groups from the preceding four hundred generations. The relatively low AMBI value in the perfect and preservation only DARCs compared to the other DARCs for the environmental change scenario reflect the strength of this “taphonomic inertia” (Kidwell 2008) with uniform time averaging (Table 3).

Differing from richness and diversity, where the most optimal values observed in any reference condition were in the perfect DARC, the optimal AMBI value was observed in the perfect LARC for both environmental scenarios. There were effectively five hundred “chances” for a generation of the live community to have a lower than average AMBI value but these short-term fluctuations were smoothed out in the time-averaged death assemblage samples (Kowalewski et al. 1998). Even so, the collective effect of using the perfect DARC rather than the perfect LARC was typically a reduction in the M-AMBI value, as the combined sensitivity of the M-AMBI values to richness and diversity often exceeded the sensitivity to AMBI values (Table 5).

The sensitivity of M-AMBI values to changes in the parameter values was variable between environmental scenarios (Table 5). When the environment was constant, M-AMBI values were most sensitive to differences in richness between the perfect DARC and the two LARCs (perfect and ten generation), followed by AMBI and then diversity (Table 5). The effects of the standardized differences for all three parameters were greater when comparing to the ten generation LARC, likely reflecting the larger overall differences in the parameters themselves (Table 3). In the environmental change scenario, differences in diversity were most influential when comparing M-AMBI results based on the perfect DARC to those that used the perfect LARC and, although the effect size for changes in diversity was larger in the ten generation LARC, it was exceeded in magnitude by the effect size for changes in AMBI values (Table 5). Richness had the smallest effect of the three reference condition parameters in both comparisons. This situational variability in the sensitivity of M-AMBI values to the three parameters highlights the importance of multivariate approaches that can integrate information from multiple metrics (Muxika et al. 2007).

#### Preservational Bias

M-AMBI scores calculated with the preservation DARC were similar to those calculated using the perfect DARC, as were the corresponding remediation decisions (Tables 2 and 4). There was a significant difference in the performance of the preservation and perfect DARCs for the constant environment simulation when compared to the perfect LARC. Even so, the difference in M-AMBI values was 0.012 (Table 3), or 1.2% of the range of possible M-AMBI values, and not likely biologically meaningful. More importantly, recall, precision, and F1 score for the preservation and perfect DARCs were nearly identical (Table 2). The largest apparent difference is in the environmental change simulations, where recall was 0.33 for the perfect DARC and 0.67 for the preservation DARC when the perfect LARC identified a need for remediation. There were, however, only three such simulation runs, meaning the apparent difference in the two DARCs amounts to variation in one out of one hundred simulation runs. Overall, the introduction of preservational bias had only minor effects compared to the perfect DARC. Given that preservational bias was random with respect to ecology in this simulation, this lack of difference is unsurprising.

Likewise, there were minor differences in the parameter values (Table 3) and the sensitivity of M-AMBI values to them (Table 5). The inclusion of preservational bias in the creation of the death assemblage led to lower probabilities that species with low preservation potential were included in the death assemblage samples. By extension, richness and diversity values in the death assemblage samples were lower than when using the perfect death assemblage (Table 3). Within environmental scenarios and at the same k values, this pattern also applies in the majority (10 of 12; Table 3) of comparisons of richness and diversity values between DA Scenario 3 (time averaging only) and DA Scenario 4 (preservational bias and time averaging). A similar effect was not observed for AMBI values because they were not affected by the distribution of species, only the distribution of ecological groups, which were random in relation to preservational bias. Though preservational bias has the potential to affect M-AMBI values, we are not currently aware of any evidence of a correlation between preservational potential and the ecological group of a species (Smith et al. 2020).

As treated here, preservational bias had a small effect on values from death assemblages but it was effectively indistinguishable from effects expected based solely on the respective time averaging scenarios. Consequently, we discuss the outcomes of DA Scenarios 3 and 4 in tandem in the next section and focus explicitly on the effects of time averaging.

#### Time averaging

Time averaging in death assemblages can be advantageous when using M-AMBI to evaluate whether a living community requires remediation. In the constant environment simulation, M-AMBI using the time-averaged death assemblages as reference conditions faithfully recapitulated all remediation decisions that were produced using M-AMBI results based on the perfect LARC. The virtues of time averaging were even more pronounced for the changing environment simulation, in which all eight of the time-averaged DARCs outperformed the ten generation LARC that was analogous to ecological monitoring.

The constant environment was overly simple and unrealistic given the prevalence of habitat change (e.g., Halpern et al. 2008). Just as the perfect LARC and perfect DARC were used to evaluate their more realistic counterparts, the constant environment simulations provided a contrast for evaluation of the changing environment simulation, particularly with respect to the effects of time averaging. In the constant simulations with variations in time averaging (DA Scenarios 3 and 4), richness and diversity values increased with time averaging as expected (Olszewski and Kidwell 2007; Kidwell 2009; Tomašovỳch and Kidwell 2010*a*) and exceeded observed values in ten and five hundred generations of the living community (Table 3). The proportional increases were relatively low (richness max = 1.29; diversity max = 1.10); however, this was likely a consequence of reporting only the maximum values used to establish reference conditions rather than mean values of samples or by comparing to only a single generation of the living community as is often done in live-dead assessments. Indeed, differences were larger when comparing to the ten generation LARC rather than the perfect LARC (Table 3). Regardless, the differences in these parameter values had only a small effect on calculated M-AMBI values when they were used as reference conditions in place of values from the perfect LARC (Figure 3). Furthermore, compared to the ten generation LARC, the DARCs more faithfully produced the correct outcomes based on the perfect LARC (Figure 2). Even in this simplistic scenario with an unrealistically stable environment, all of the DARCs outperformed the ten generation LARC as reference conditions.

The merits of time averaging and the effect of different durations of time averaging were more pronounced in the environmental change simulations. As time averaging decreased (i.e., k- values increased), the differences in parameter values and the corresponding M-AMBI values increased relative to the perfect DARC (Table 3). That is, the parameter values used to define the reference conditions became less optimal and by comparison, the ecological quality of the living community appeared less degraded. Compared to the perfect LARC, use of DARCs with little time averaging (i.e., k= 1.5) led to overestimation of M-AMBI values by as much as 0.088, or nearly 10% the possible range of M-AMBI values (Table 3). This overestimation corresponded to the increasing proportion of individuals in the death assemblages that lived after the environmental change. When the k-value was low (i.e., 0.5), time averaging spanned 495 generations on average and the mean age of an individual in the death assemblage was 171 - 173 (i.e., died at generation 329 - 327). At the other extreme (k = 1.5), time averaging encompassed an average of 296 - 451 generations and the mean age of an individual was 67 - 80. At all k-values, the age-frequency distributions were skewed towards younger generations in DA Scenarios 3 and 4 (see supplementary materials Table S3) but this pull of the Recent only became pronounced at low levels of time averaging.

As the extent of time averaging in the death assemblage decreased, the differences in M-AMBI values calculated using DARCs and the perfect LARC increased (Table 3). This trend in M-AMBI values was directly related to corresponding changes to the parameter values, as all values became less optimal: richness and diversity values decreased and AMBI values increased (Table 3). There was a similar trend in values when the environment was constant, however, the magnitude of differences in the range of values in the environmental change simulations was far greater (Table 3). For example, the difference in DARC richness between the highest and lowest k-values was more than five times larger in the environmental change scenario (7.68 vs. 1.38; Low TA w/ pres). The effect of decreasing time averaging was compounded by the environmental change. Because mollusks are relatively sensitive to disturbance, when the environment shifted to a disturbed, lower quality condition, the number of species capable of surviving (defined by the ecological trait) decreased and the death assemblage became dominated by relatively few, opportunistic species – a phenomenon that is also observed naturally (e.g., *Mulinia*; Levinton 1970; Shumway and Newell 1984; Kowalewski et al. 2000; Dietl and Smith 2017). The maximum richness values in the perfect LARC and DARCs with less time averaging (k = 1.5) were nearly four times greater than the ten generation LARC, and DARCs with more time averaging had even larger differences. Likewise, maximum diversity in the ten generation LARC was approximately half of the values observed for the other reference conditions. These patterns in richness and diversity align well with patterns observed in the literature, demonstrating that increased time averaging can lead to increases in richness and diversity (e.g., Olszewski and Kidwell 2007; Tomašovỳch and Kidwell 2010; Bürkli and Wilson 2017).

Time averaging also affected the sensitivity of M-AMBI values to changes in the DARC parameter values in simulations with environmental change. Relative to the perfect LARC, differences in the diversity values used to calculate M-AMBI had the largest influence. This influence diminished as time averaging was reduced to the point where the effect sizes of all three parameter values were similar (Table 5). By contrast, the effect of changes in richness and AMBI increased slightly as time averaging decreased. As time averaging decreased, richness values in the DARCs approached the values from the perfect LARC but were always more optimal. Similarly, AMBI values in the perfect LARC were always more optimal than in the DARCs. The most optimal diversity values switched from the DARC to the perfect LARC as time averaging decreased and the death assemblage became dominated by species from post-environmental change generations, likely explaining the pattern in M-AMBI sensitivity to changes in parameter values. The pattern was simpler in comparisons between the DARCs and the ten generation LARC as the effect of diversity behaved like richness and AMBI: effect size increased as time averaging decreased and the values observed in the DARCs were consistently more optimal by large margins. The effect size of parameter changes was largest for AMBI in these comparisons with the ten generation LARC, rather than diversity (as in comparison to the perfect LARC) or richness (as in the constant environment simulations), emphasizing the importance of considering more than a single parameter (Muxika et al. 2007). Likewise, the sensitivity of parameter and M-AMBI values to time averaging in the death assemblages underscores the importance of quantifying time averaging when drawing ecological conclusions (Kidwell 2013; Kidwell and Tomašovỳch 2013).

### Shifting baselines

In the simulation runs with and without environmental change, M-AMBI values calculated using the ten generation LARC deviated substantially from the values calculated using the perfect LARC. The simulation was constructed in such a way that this result was to be expected. Even though the ten generations and five hundred generations in the constant environment simulation runs used the same environmental filter, the number of generations used to calculate the reference conditions all but guaranteed that more optimal values would be found with the longer option due to chance. Similarly, by the introduction of the environmental change one hundred generations prior to the end of the simulation and limitation of the LARC to only the final ten generations, it was inevitable that the ten generation LARC would be biased compared to the perfect LARC (Figure 2). Nonetheless, these simulations demonstrate the potential perils of using a limited temporal perspective to establish reference conditions (Pauly 1995). Considering the limited duration of many ecological monitoring efforts and the challenges of maintaining such efforts (Lindenmayer et al. 2012), it can be challenging to avoid the effects of shifting baseline syndrome.

Death assemblages provide a means to safeguard against shifting baseline syndrome when the extent of time averaging in a death assemblage exceeds the time since environmental change. The simulation used here provides proof of concept. With and without environmental change, all instances with a death assemblage based reference condition produced more faithful estimates of M-AMBI than the ten generation LARC (Figure 2, Table 3). Death assemblages can also be used to detect improvements to ecological quality, potentially reflecting remediation or policy outcomes. For example, in Long Island Sound comparisons of living and death assemblages to evaluate the effects of hypoxia and fishing pressure revealed an unexpected improvement in ecological quality of habitats closed to fishing (Casey et al. 2014). In this way, death assemblages can be used not only to establish reference conditions but also to evaluate the outcomes of past actions.

### Limitations and future work

The simulation framework developed here relied on several simplifying assumptions and does not incorporate all of the potential biases that may affect death assemblages. Even so, the framework is parameterized by real ecological observations and expands on the ecological theory used in previous simulations (e.g., neutral theory, density-dependence, dispersal limitation, life-history trade-offs; (Tomašovỳch and Kidwell 2010*a*, *b*; Tomašových and Kidwell 2011)) through the incorporation of environmental filtering, which has been shown to contribute to molluscan community assembly (Moritz et al. 2009; Noda 2009; Valdivia et al. 2015; Smith and Dietl 2019). The richness and diversity values from the living assemblages and their differences with values from the various death assemblage scenarios also compare well to real observations of live-dead differences (e.g., Kidwell 2002, 2007; Olszewski and Kidwell 2007; Tomašovỳch and Kidwell 2010; Bürkli and Wilson 2017). As such, the simulation provides an acceptable approximation of reality. Expansion of the simulation framework through testing of assumptions and incorporation of additional biases will allow for stronger insights into ecological and taphonomic processes and patterns recoverable from death assemblages.

This simulation approach is based on several simplifying assumptions used to reduce the complexity of ecological and preservation processes. In particular, the effects of ecology and preservation are each represented by a single highly generalized variable. We use these simplified variables to incorporate the importance of ecological and preservational processes without becoming overburdened by trying to evaluate their intricacies (Barido-Sottani et al. 2020), as the intention here is primarily to consider the effects of time averaging on M-AMBI values when using death assemblages as reference conditions. Future studies might evaluate the effect of incorporating multiple ecological and preservational traits, and allowing correlations within and between them. As in most simulations, we assumed that many parameters remained constant and allowing variation in any of these would prove fruitful for future study. Among those assumptions were: (i) a constant, non-evolving metacommunity, (ii) equivalent dispersal capacity of all species, (iii) no competition between species in the local community, (iv) constant community size, (v) equal life spans of all species, and (vi) transference of results of a spatially implicit model to a three-dimensional space. In future work, the simulation framework developed here can be used without one or more of these assumptions to evaluate other patterns and processes.

In addition, as our focus was on the effects of time averaging, several other biases were not evaluated. These include (i) life span bias; (ii) proportional abundance of mollusks in the living community; and, (iii) molluscan sensitivity to disturbance. Though not incorporated here, the simulation framework has the capacity to incorporate these factors in future iterations. Life span bias may be particularly important because differences in population turnover rates in living communities have the potential to bias species abundances observed in death assemblages (Cronin et al. 2018). Species with faster rates of turnover will produce more shells per annum than species with slower rates. Consequently, short-lived species may be overrepresented in death assemblages compared to longer lived species (Cronin et al. 2018). Because opportunistic species also tend to be shorter lived, life span bias may lead to artificially high AMBI values in the death assemblage. Reference conditions from such a death assemblage are expected to be less optimal and EcoQS of the living communities would likely be overestimated. Using the simulation to adjust species-specific death rates may help evaluate the magnitude of this potential bias.

There may also be an interaction between life span bias and the type of disturbance to the environment. The disturbance simulated here caused an abrupt and permanent change–as the environmental filter switched from 0.2 at generation 400 to 0.8 at generation 401–and was akin to rapid changes experienced during a regime shift (deYoung et al. 2008). A disturbance may instead be abrupt but temporary (i.e., pulse), be a permanent monotonic shift (i.e., press; (Bender et al. 1984), or intermediates of these two options driven by variability in the durations over which the causes and effects of the disturbance persist (Glasby and Underwood 1996). Short- lived, opportunistic species thrive during periods of disturbance and a death assemblage may accumulate a large influx of such species (Levinton 1970; Levinton and Bambach 1970), which will only increase as the duration of a disturbance increases. A greater breadth of disturbance patterns can be incorporated into the simulation framework, with and without life span bias, to evaluate the effects of these features in isolation and in combination.

The other two potential biases discussed by Smith et al. (2020), proportional abundance of mollusks and molluscan sensitivity to disturbance, can be evaluated by changing the framing of the simulation. Because death assemblages were the focus here, the set of species that were simulated were all considered mollusks *a priori*. By increasing the number of species and the size of the metacommunity, the simulation can accommodate the inclusion of species considered to belong to other phyla (e.g., arthropods, foraminifera, worms). The distribution of these taxa and associated ecological groups can be assigned in the same manner as used for mollusks here. Inclusion of this broader taxonomic set would also allow for examination of the assumption used here that M-AMBI scores calculated for mollusks have a 1:1 mapping to the entire benthic community. Equations for correcting mollusk-only M-AMBI scores to represent the entire community can be drawn from Dietl et al. (2016) and Pruden et al. (2021), and these equations can themselves be evaluated with the simulation framework. Evaluating these biases will be important because the capacity of the mollusk-only index to recapitulate values from the whole-community index weakens as the proportion of mollusks in the community decreases (Dietl et al. 2016). Similarly, the majority of molluscan species are categorized in low ecological groups, meaning they are more sensitive to disturbance than other taxonomic groups. Consequently, mollusks are often absent from highly disturbed sites and the effectiveness of mollusks as sensors in highly disturbed environments may be low.

The assumptions and biases discussed in this section have the potential to alter the M-AMBI scores observed when using reference conditions based on death assemblages. Further study is needed to evaluate the magnitude and direction of these effects on M-AMBI scores, and the interactions between effects. Though more work is required, it is encouraging that the major and most prevalent feature of death assemblages, time averaging, is an advantage rather than a detriment. Through quantification of these biases and verification of the results with real-world case studies, conservation paleobiology can contribute to the management of estuarine ecosystems by filling a need for improved reference conditions.

### Conclusion

The simulations evaluated here suggest that death assemblages can perform well as reference conditions for M-AMBI. In the simulations with a constant environment, reference conditions from all eight versions of the death assemblage perfectly reproduced the remediation decisions and EcoQS produced when the M-AMBI reference conditions were based on perfect knowledge of the entire five hundred generations in the simulation. When the environment changed, the death assemblage reference conditions continued to perform well and reproduced the “correct” result in at least 75% of simulation runs. Time averaging was a beneficial feature of death assemblages, as they retained a memory of pre-change generations of the community. By comparison, use of reference conditions based on only the ten most recent generations of the simulation succumbed to shifting baseline syndrome, overestimating the quality of the living community and failing to indicate a need for remediation when dictated by the perfect knowledge reference condition.

Increased richness and diversity in death assemblages affected M-AMBI scores as predicted. By increasing the ecological quality implied by the reference conditions through increased richness and diversity, use of death assemblages as reference conditions with increasing values (a consequence of increased time averaging), led to lower M-AMBI values in the living community. Even so, the M-AMBI values were consistently similar to those calculated using the perfect knowledge reference conditions. That is, the “true” optimal values for richness, diversity, and AMBI based on all five hundred generations were similar to those observed in the death assemblages. The sensitivity of M-AMBI values to changes in the different parameters was variable between the two simulations, but overall the simulations suggest that death assemblages produce values that can be reliably used to generate reference conditions. Based on these simulations, we conclude: (1) it is important to evaluate information from multiple metrics together, for example, with multivariate approaches ; (2) studies using death assemblages need to evaluate the duration of time averaging; and, (3) particularly when the duration of time averaging is known, death assemblages can provide valuable long term perspectives with the potential to outperform temporally constrained baseline information from live observations.

## Author contributions

**Jansen A. Smith:** Conceptualization, Investigation, Methodology, Software, Writing - original draft, Writing - review & editing; **Matthew J. Pruden:** Conceptualization, Formal analysis, Methodology, Software, Visualization, Writing - original draft, Writing - review & editing; **John C. Handley:** Conceptualization, Formal analysis, Methodology, Writing - review & editing; **Stephen R. Durham:** Conceptualization, Writing - review & editing; **Gregory P. Dietl:** Conceptualization, Writing - review & editing

## SUPPLEMENTARY MATERIAL

### Expanded history of biotic indices

The composition of aquatic ecological communities is strongly linked to environmental conditions, and therefore can provide information on the effects of water pollution (De Pauw and Vanhooren 1983; Metcalfe 1989; Kröncke and Reiss 2010; Rousi et al. 2019; Carrier-Belleau et al. 2021). Benthic macroinvertebrates are easy to collect, differentially sensitive to various physio-chemical conditions, and are relatively sedentary and therefore representative of local conditions (Cook 1976; Hily 1984; Dauer 1993; Diaz and Rosenberg 1995; Lamouroux et al. 2004; McCormick et al. 2004). While benthic macroinvertebrate biotic indices originated in the United States (e.g., Gaufin and Tarzwell 1952, 1956), it was not until the development of the Trent Biotic Index (TBI; Woodiwiss 1964) for freshwater systems in England that benthic indices began to grow in prominence. Woodiwiss (1964) was one of the first to propose a biotic index directly linking changes in benthic community composition to organic enrichment, which has since formed the basis for most modern biotic indices (Persoone and De Pauw 1979; Metcalfe 1989). The TBI is based on the presence of organisms from key groups and the number of groups in a sample, with each group having a defined sensitivity to organic enrichment (the groups are assigned at either the Family, Genus, or species level depending on the type of organism; Woodiwiss 1964). Though the TBI and its derivatives showed promise for freshwater systems, it was unknown if such models could be extended to the marine realm, due to the uncertainty in the response of marine benthic macroinvertebrate communities to organic enrichment given the greater variety of habitats and species (Bellan and Bellan-Santini 1972; Reish 1972; Pearson and Rosenberg 1978).

Pearson and Rosenberg (1978) sought to alleviate the uncertainty and synthesized the existing knowledge of the effects of organic enrichment on the composition of benthic macroinvertebrate communities within the marine realm. They hypothesized that macroinvertebrate species could be divided into two general groups based on their physiological characteristics and response to enrichment. In areas of high organic enrichment, the community is often composed of short-lived, small macroinvertebrates with high reproductive rates. As the degree of enrichment decreases, the composition of the community changes to include a wider range of ecological and reproductive characteristics (Pearson and Rosenberg 1978). Gray (1979) expanded Pearson and Rosenberg’s (1978) groupings by dividing benthic macroinvertebrate species into three ecological groups based on their reproductive strategy and physiological adaptations to organic enrichment: (i) k-selected species, species with relatively long-life spans, slow growth, and adapted to live in areas with low physical disturbance (examples of physical disturbances include: storms, smothering, high wave energy, and dredging); (ii) r-selected species, species with short-life spans, fast growth, and adapted to live under high physical disturbance; and, (iii) T-stress tolerant species, species which are adapted to live under low physical disturbance but high environmental stress (e.g., low salinity, high organic enrichment).

Finally, Hily (1984), Glemarec (1986), and Grall and Gelmarec (1997) divided the macroinvertebrate species further into five ecological groups (EGs) of increasing tolerance to organic enrichment- EGI = species sensitive to organic enrichment; EGII = species indifferent to enrichment and always present in low densities; EGIII = species tolerant to enrichment and whose populations are stimulated by organic enrichment; EGIV and EGV = second- and first- order opportunistic species, respectively, with short life cycles and adaptations to life in reduced sediments. Note, the number of ecological groups is not an indicator of index precision, nor are the ecological groupings discussed herein the only viable classifications. There are other widely used biotic indices that utilize fewer or more groupings than are summarized in Grall and Glemarec (1997) (e.g., BENTIX; Simboura and Zenetos 2002). We focus on the development of AMBI and M-AMBI, hence our selective discussion of the groupings relevant to these indices.

### Simulation elements

**Figure S1.**
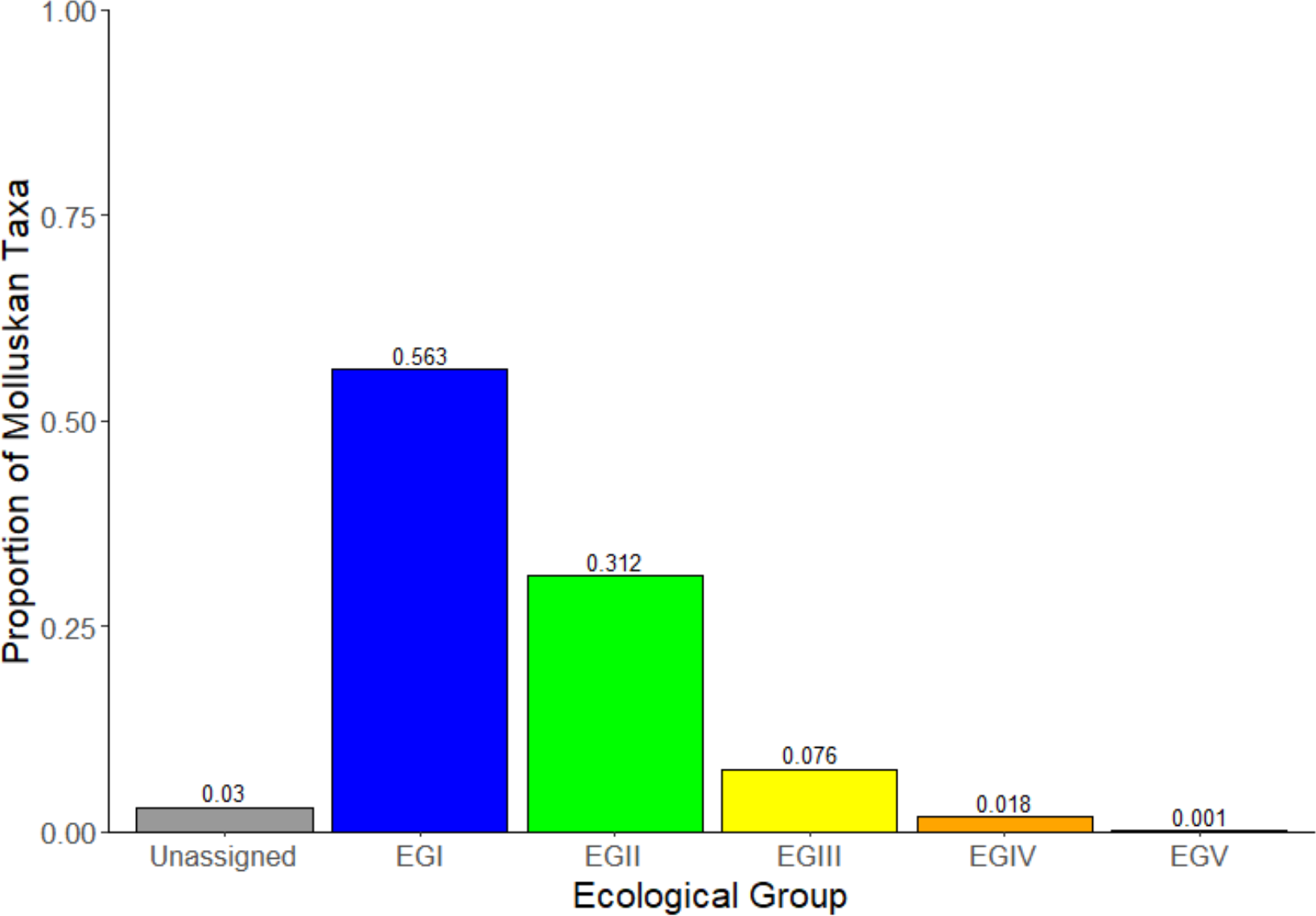
Distribution of molluscan species in the AMBI software v6.0 (http://ambi.azti.es; Species List v.Dec2020)

**Figure S2.**
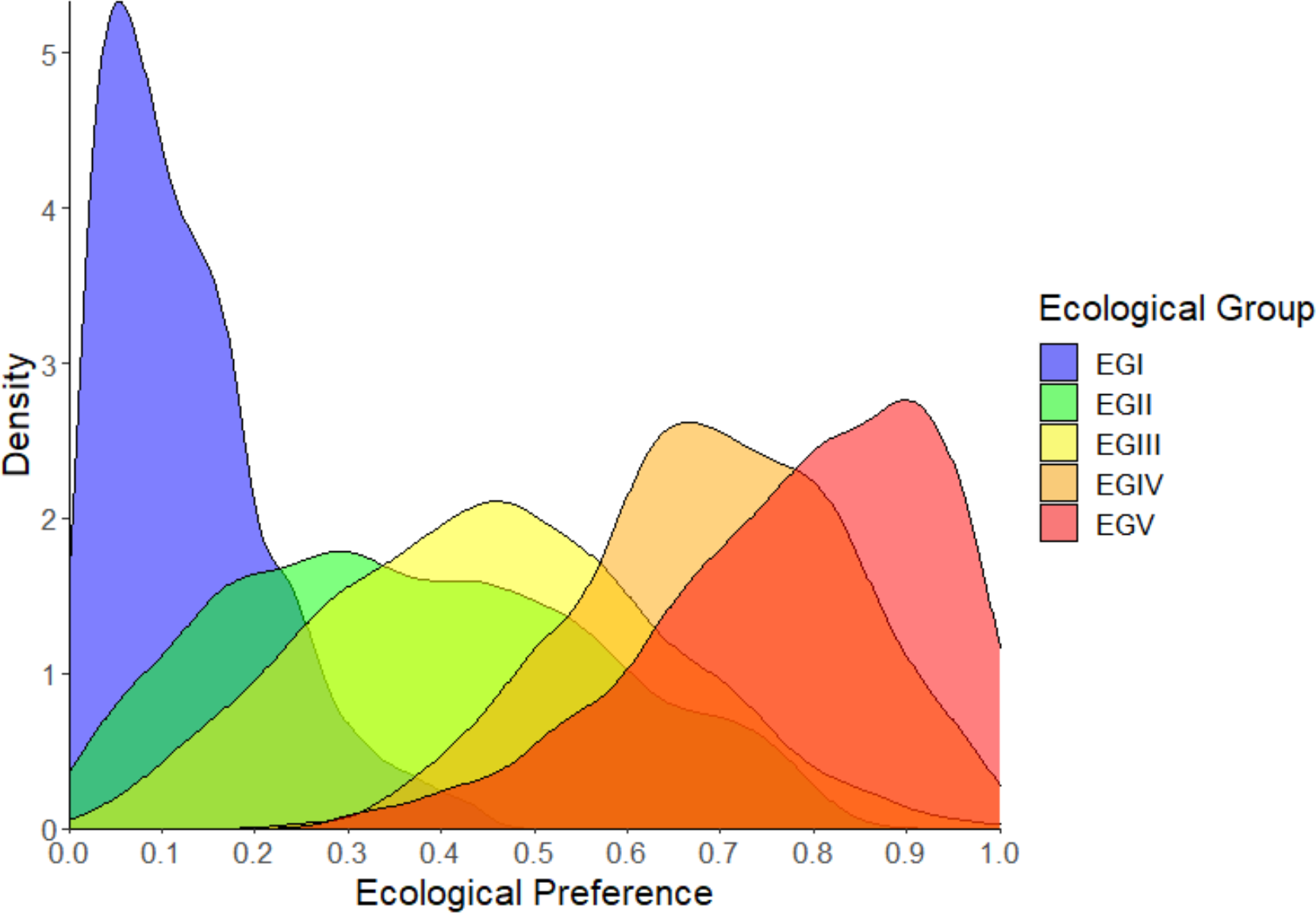
Distributions of molluscan ecological trait values in the five ecological groups used to calculate AMBI (Borja et al. 2000) and to parameterize the simulation.

### Statistical decisions

The non-parametric Kruskall-Wallis test, with the Holm p-value adjustment method to control the familywise error rate, was used because the assumption of normality was not upheld in all comparisons (Aickin and Gensler 1996; Hecke 2012). See figures S3 - S6 for specifics. Constant Env = Constant environment scenario; Env Change = Environmental change scenario; Perfect LARC = All 500 Generation Living Assemblage Reference Condition; 10 Gen LARC = 10 Generation Living Assemblage Reference Condition; DARC = Death Assemblage Reference Condition; TA = Time-Averaging; Pres = Differential Preservation.

**Figure S3.**
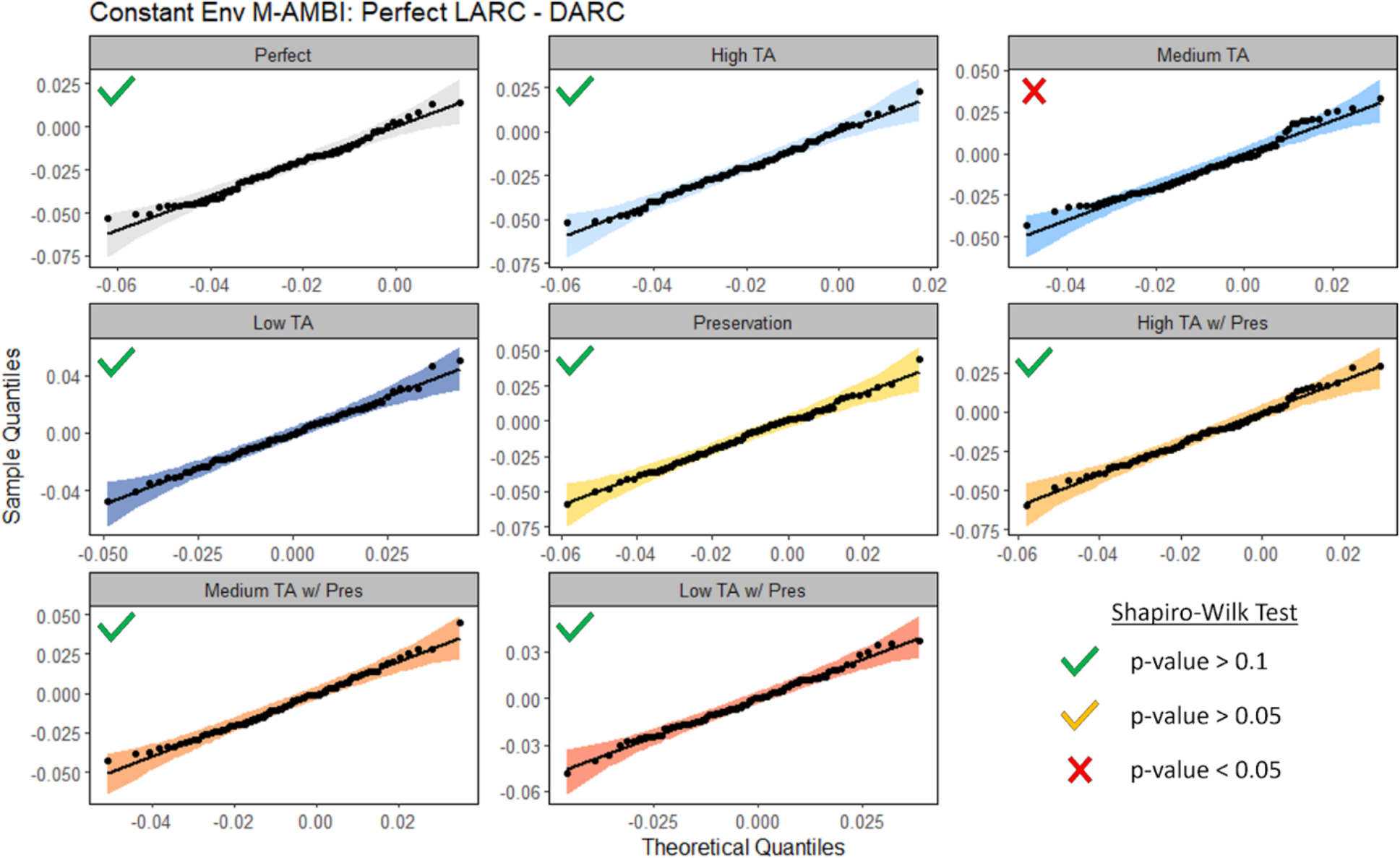
Quantile-quantile plots to evaluate normality in comparisons between the perfect live assemblage reference condition and the various death assemblage reference conditions in the simulations with a constant environment. LARC = Live assemblage reference condition; DARC = Death assemblage reference condition; TA = time averaging; w/Pres indicates the inclusion of between species differences in preservation potential.

**Figure S4.**
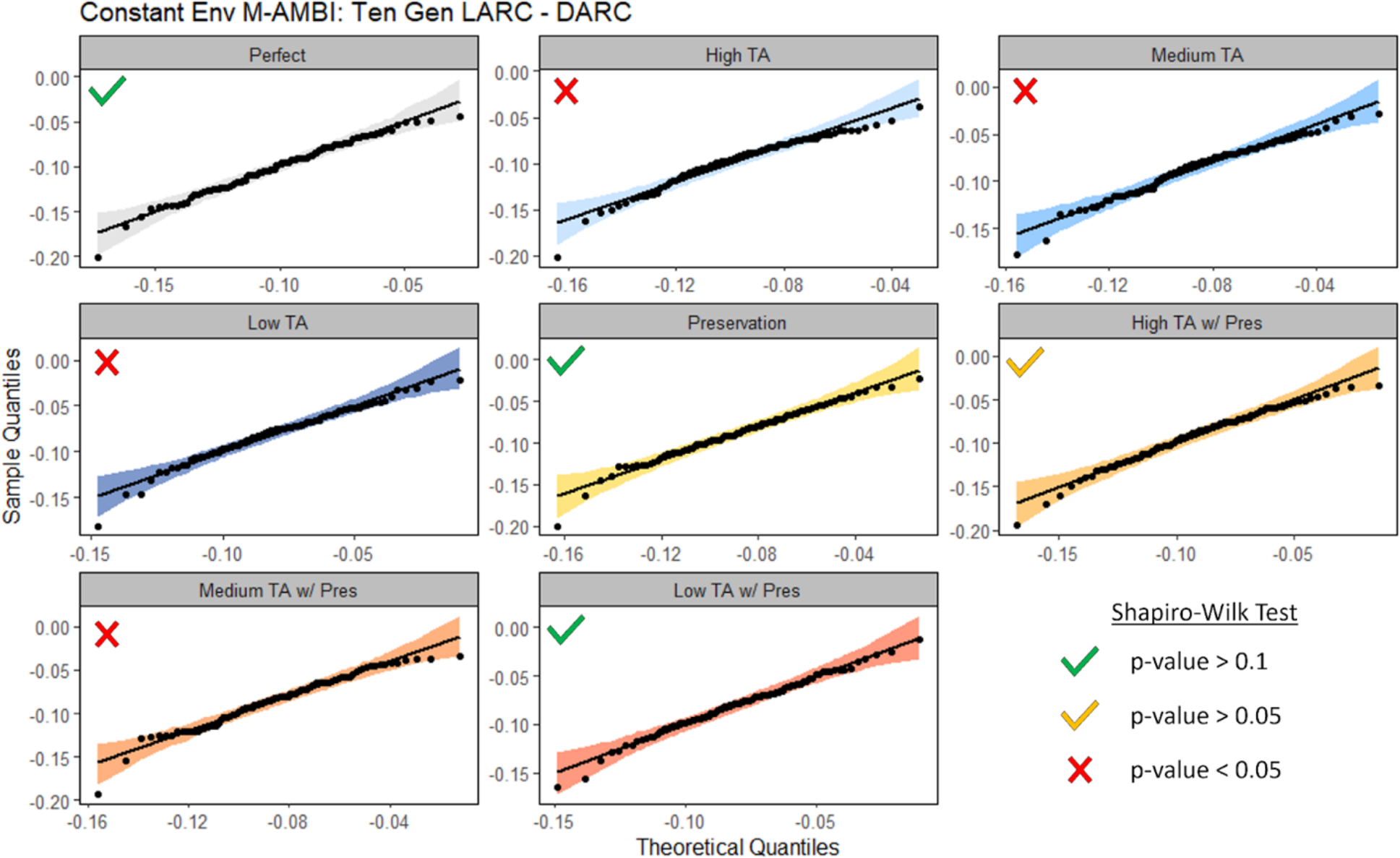
Quantile-quantile plots to evaluate normality in comparisons between the ten generation live assemblage reference condition and the various death assemblage reference conditions in the simulations with a constant environment. LARC = Live assemblage reference condition; DARC = Death assemblage reference condition; TA = time averaging; w/Pres indicates the inclusion of between species differences in preservation potential.

**Figure S5.**
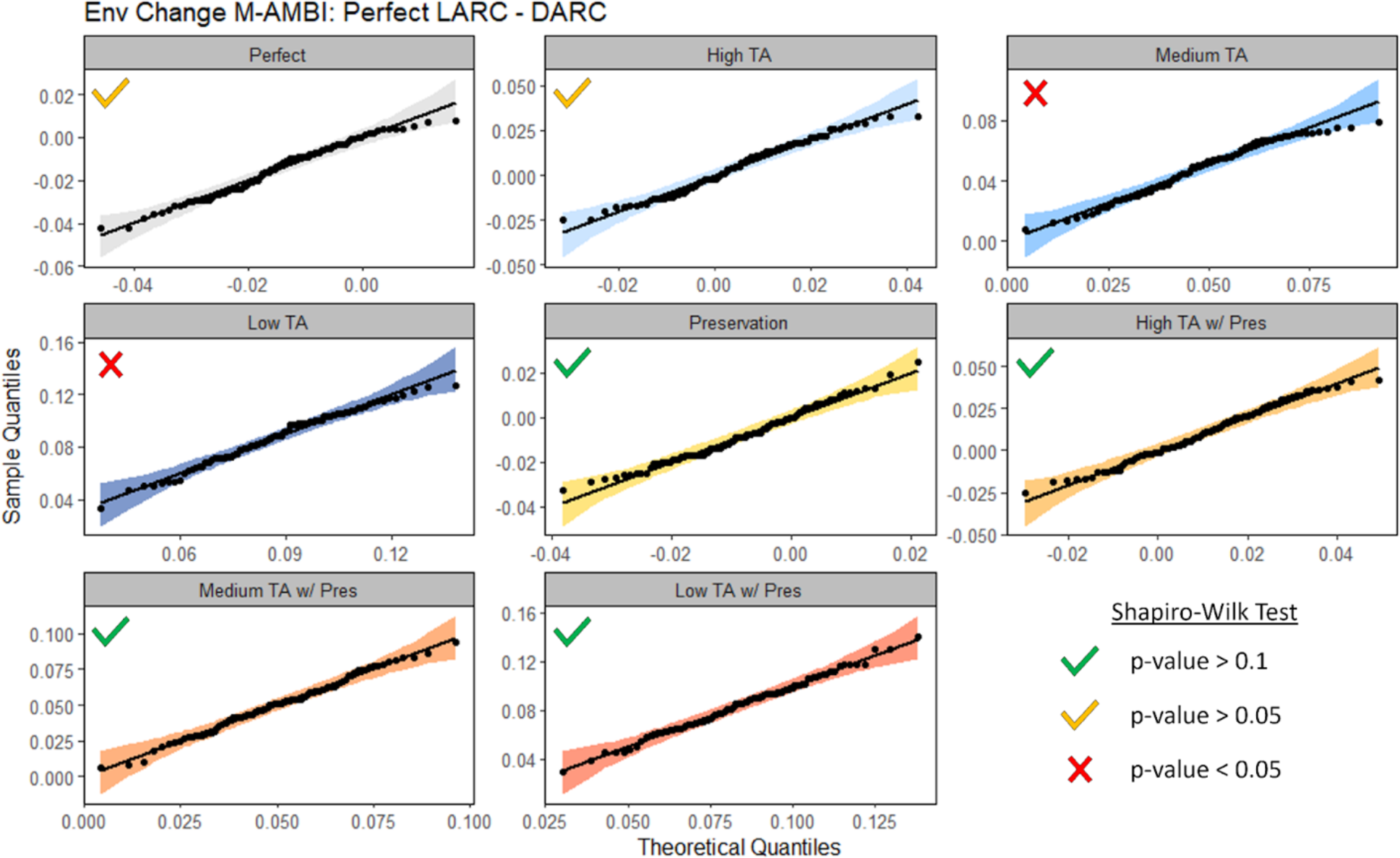
Quantile-quantile plots to evaluate normality in comparisons between the perfect live assemblage reference condition and the various death assemblage reference conditions in the simulations with environmental change. LARC = Live assemblage reference condition; DARC = Death assemblage reference condition; TA = time averaging; w/Pres indicates the inclusion of between species differences in preservation potential.

**Figure S6.**
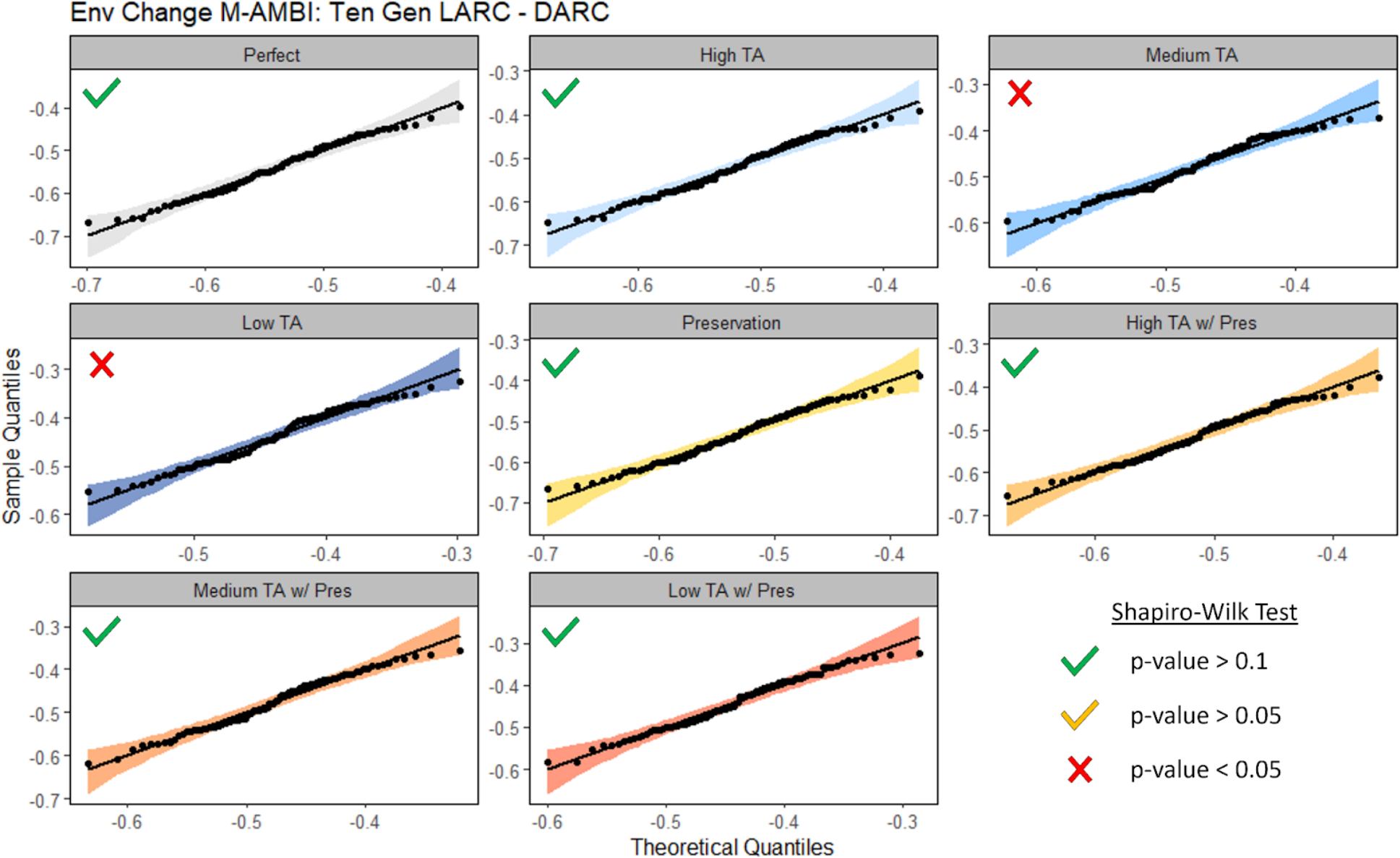
Quantile-quantile plots to evaluate normality in comparisons between the ten generation live assemblage reference condition and the various death assemblage reference conditions in the simulations with environmental change. LARC = Live assemblage reference condition; DARC = Death assemblage reference condition; TA = time averaging; w/Pres indicates the inclusion of between species differences in preservation potential.

### Ancillary simulation outputs

**Table S1.**
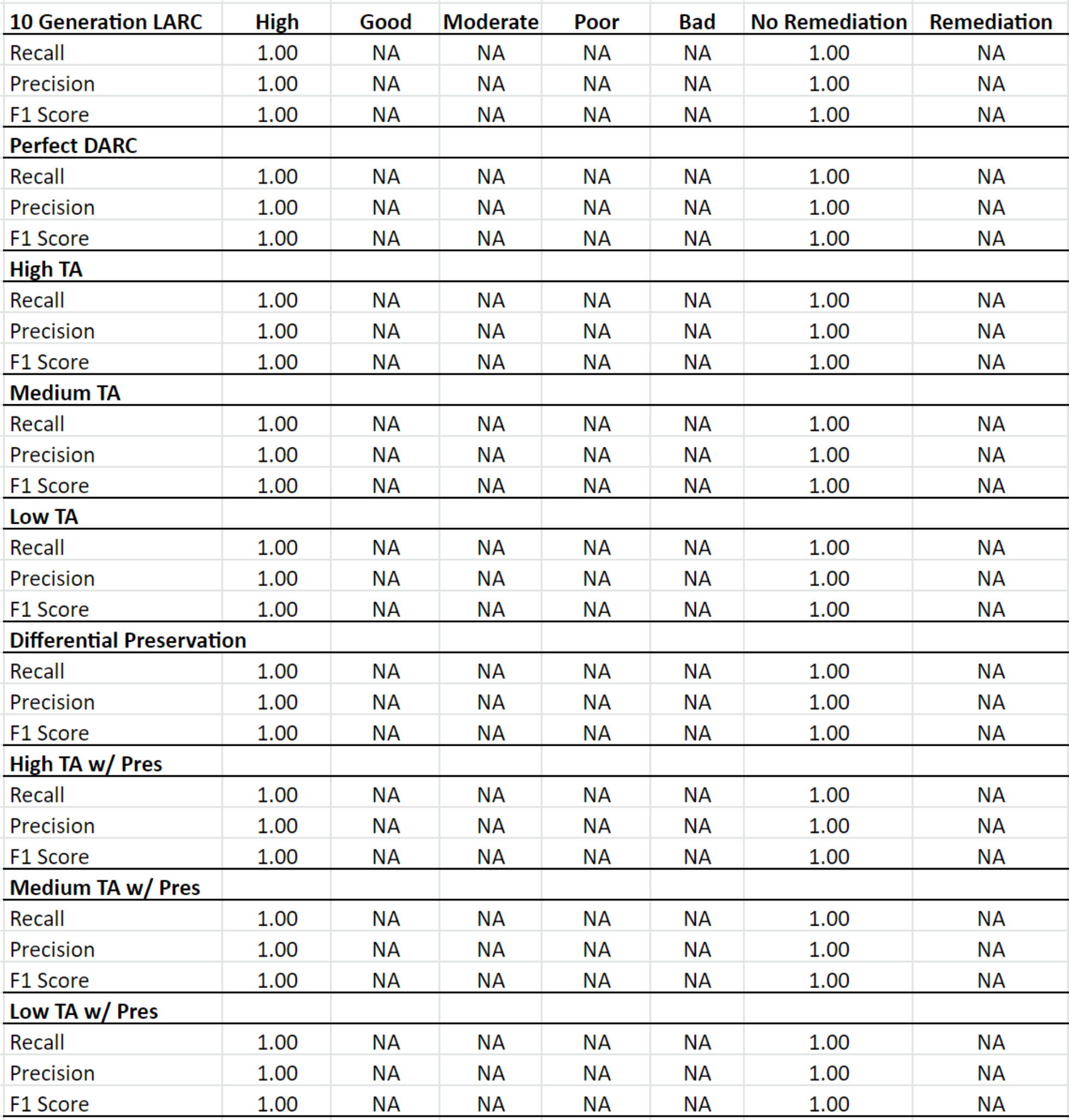
Fidelity metric scores for all Ecological Quality Statuses in simulations with a constant environment. LARC = Live assemblage reference condition; DARC = Death assemblage reference condition; TA = time averaging; w/Pres indicates the inclusion of between species differences in preservation potential.

| **10 Generation LARC** | **High** | **Good** | **Moderate** | **Poor** | **Bad** | **No Remediation** | **Remediation** |
| --- | --- | --- | --- | --- | --- | --- | --- |
| Recall | 1.00 | NA | NA | NA | NA | 1.00 | NA |
| Precision | 1.00 | NA | NA | NA | NA | 1.00 | NA |
| F1 Score | 1.00 | NA | NA | NA | NA | 1.00 | NA |
| **Perfect DARC** |  |  |  |  |  |  |  |
| Recall | 1.00 | NA | NA | NA | NA | 1.00 | NA |
| Precision | 1.00 | NA | NA | NA | NA | 1.00 | NA |
| F1 Score | 1.00 | NA | NA | NA | NA | 1.00 | NA |
| **High TA** |  |  |  |  |  |  |  |
| Recall | 1.00 | NA | NA | NA | NA | 1.00 | NA |
| Precision | 1.00 | NA | NA | NA | NA | 1.00 | NA |
| F1 Score | 1.00 | NA | NA | NA | NA | 1.00 | NA |
| **Medium TA** |  |  |  |  |  |  |  |
| Recall | 1.00 | NA | NA | NA | NA | 1.00 | NA |
| Precision | 1.00 | NA | NA | NA | NA | 1.00 | NA |
| F1 Score | 1.00 | NA | NA | NA | NA | 1.00 | NA |
| **Low TA** |  |  |  |  |  |  |  |
| Recall | 1.00 | NA | NA | NA | NA | 1.00 | NA |
| Precision | 1.00 | NA | NA | NA | NA | 1.00 | NA |
| F1 Score | 1.00 | NA | NA | NA | NA | 1.00 | NA |
| **Differential Preservation** |  |  |  |  |  |  |  |
| Recall | 1.00 | NA | NA | NA | NA | 1.00 | NA |
| Precision | 1.00 | NA | NA | NA | NA | 1.00 | NA |
| F1 Score | 1.00 | NA | NA | NA | NA | 1.00 | NA |
| **High TA w/ Pres** |  |  |  |  |  |  |  |
| Recall | 1.00 | NA | NA | NA | NA | 1.00 | NA |
| Precision | 1.00 | NA | NA | NA | NA | 1.00 | NA |
| F1 Score | 1.00 | NA | NA | NA | NA | 1.00 | NA |
| **Medium TA w/ Pres** |  |  |  |  |  |  |  |
| Recall | 1.00 | NA | NA | NA | NA | 1.00 | NA |
| Precision | 1.00 | NA | NA | NA | NA | 1.00 | NA |
| F1 Score | 1.00 | NA | NA | NA | NA | 1.00 | NA |
| **Low TA w/ Pres** |  |  |  |  |  |  |  |
| Recall | 1.00 | NA | NA | NA | NA | 1.00 | NA |
| Precision | 1.00 | NA | NA | NA | NA | 1.00 | NA |
| F1 Score | 1.00 | NA | NA | NA | NA | 1.00 | NA |

**Table S2.** Fidelity metric scores for all Ecological Quality Statuses in simulations with environmental change. LARC = Live assemblage reference condition; DARC = Death assemblage reference condition; TA = time averaging; w/Pres indicates the inclusion of between species differences in preservation potential.

| **10 Generation LARC** | **High** | **Good** | **Moderate** | **Poor** | **Bad** | **No Remediation** | **Remediation** |
| --- | --- | --- | --- | --- | --- | --- | --- |
| Recall | NA | 0.00 | 0.00 | 0.00 | NA | 1.00 | 0.00 |
| Precision | 0.00 | NA | NA | NA | NA | 0.03 | NA |
| F1 Score | NA | NA | NA | NA | NA | 0.06 | NA |
| **Perfect DARC** |  |  |  |  |  |  |  |
| Recall | NA | 0.33 | 0.86 | 1.00 | NA | 0.33 | 1.00 |
| Precision | NA | 1.00 | 0.96 | 0.79 | NA | 1.00 | 0.98 |
| F1 Score | NA | 0.50 | 0.91 | 0.88 | NA | 0.50 | 0.99 |
| **High TA** |  |  |  |  |  |  |  |
| Recall | NA | 0.67 | 0.97 | 0.76 | NA | 0.67 | 1.00 |
| Precision | NA | 1.00 | 0.87 | 0.93 | NA | 1.00 | 0.99 |
| F1 Score | NA | 0.80 | 0.92 | 0.84 | NA | 0.80 | 0.99 |
| **Medium TA** |  |  |  |  |  |  |  |
| Recall | NA | 1.00 | 0.95 | 0.24 | NA | 1.00 | 0.97 |
| Precision | NA | 0.50 | 0.70 | 1.00 | NA | 0.50 | 1.00 |
| F1 Score | NA | 0.67 | 0.81 | 0.38 | NA | 0.67 | 0.98 |
| **Low TA** |  |  |  |  |  |  |  |
| Recall | NA | 1.00 | 0.44 | 0.09 | NA | 1.00 | 0.64 |
| Precision | NA | 0.08 | 0.47 | 1.00 | NA | 0.08 | 1.00 |
| F1 Score | NA | 0.15 | 0.46 | 0.16 | NA | 0.15 | 0.78 |
| **Differential Preservation** |  |  |  |  |  |  |  |
| Recall | NA | 0.67 | 0.90 | 0.97 | NA | 0.67 | 1.00 |
| Precision | NA | 1.00 | 0.97 | 0.85 | NA | 1.00 | 0.99 |
| F1 Score | NA | 0.80 | 0.93 | 0.90 | NA | 0.80 | 0.99 |
| **High TA w/ Pres** |  |  |  |  |  |  |  |
| Recall | NA | 1.00 | 0.98 | 0.74 | NA | 1.00 | 1.00 |
| Precision | NA | 1.00 | 0.87 | 0.96 | NA | 1.00 | 1.00 |
| F1 Score | NA | 1.00 | 0.93 | 0.83 | NA | 1.00 | 1.00 |
| **Medium TA w/ Pres** |  |  |  |  |  |  |  |
| Recall | NA | 1.00 | 0.87 | 0.32 | NA | 1.00 | 0.92 |
| Precision | NA | 0.27 | 0.71 | 1.00 | NA | 0.27 | 1.00 |
| F1 Score | NA | 0.43 | 0.78 | 0.49 | NA | 0.43 | 0.96 |
| **Low TA w/ Pres** |  |  |  |  |  |  |  |
| Recall | NA | 1.00 | 0.48 | 0.09 | NA | 1.00 | 0.66 |
| Precision | NA | 0.08 | 0.49 | 1.00 | NA | 0.08 | 1.00 |
| F1 Score | NA | 0.15 | 0.48 | 0.16 | NA | 0.15 | 0.80 |

**Table S3.** Mean duration of time averaging and mean age of individuals across simulations with and without environmental change. LARC = Live assemblage reference condition; DARC = Death assemblage reference condition; TA = time averaging; w/Pres indicates the inclusion of between species differences in preservation potential.

| DARC | k-value | Constant Environment |  | Environmental Change |  |
| --- | --- | --- | --- | --- | --- |
|  |  | Mean Age | TA Duration | Mean Age | TA Duration |
| Perfect | NA | 250.51 | 496.01 | 250.51 | 496.01 |
| High TA | 0.5 | 173.18 | 494.58 | 173.18 | 494.58 |
| Medium TA | 1.0 | 97.39 | 457.81 | 97.39 | 457.81 |
| Low TA | 1.5 | 66.65 | 295.90 | 66.65 | 295.90 |
| Differential Preservation | NA | 250.41 | 495.91 | 249.97 | 495.89 |
| High TA w/ Pres | 0.5 | 171.96 | 494.42 | 171.44 | 494.54 |
| Medium TA w/ Pres | 1.0 | 105.30 | 478.84 | 104.64 | 478.28 |
| Low TA w/ Pres | 1.5 | 80.60 | 452.97 | 80.17 | 450.75 |

**Figure S7.**
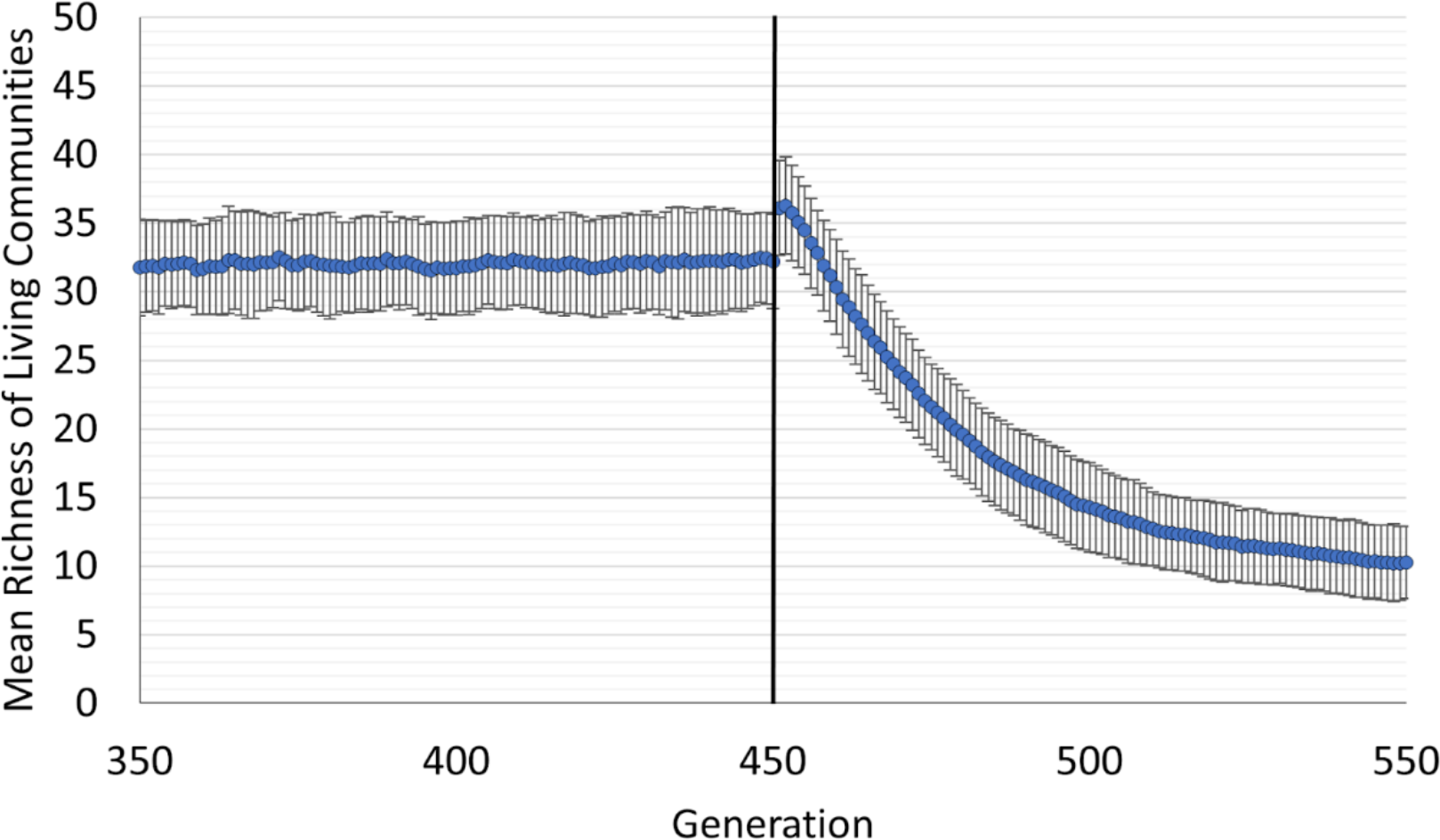
Richness in living communities 100 generations before and after environmental change, across all 100 simulations. The initial increase after generation 450 reflects persistence of the previous high ecological quality community (i.e., priority effects) coupled with the immigration of new, low quality species. Overall changes in community composition lagged behind the rapid change in environmental conditions. The black horizontal line corresponds to the timing of environmental change. Error bars represent one standard deviation.

**Figure S8.**
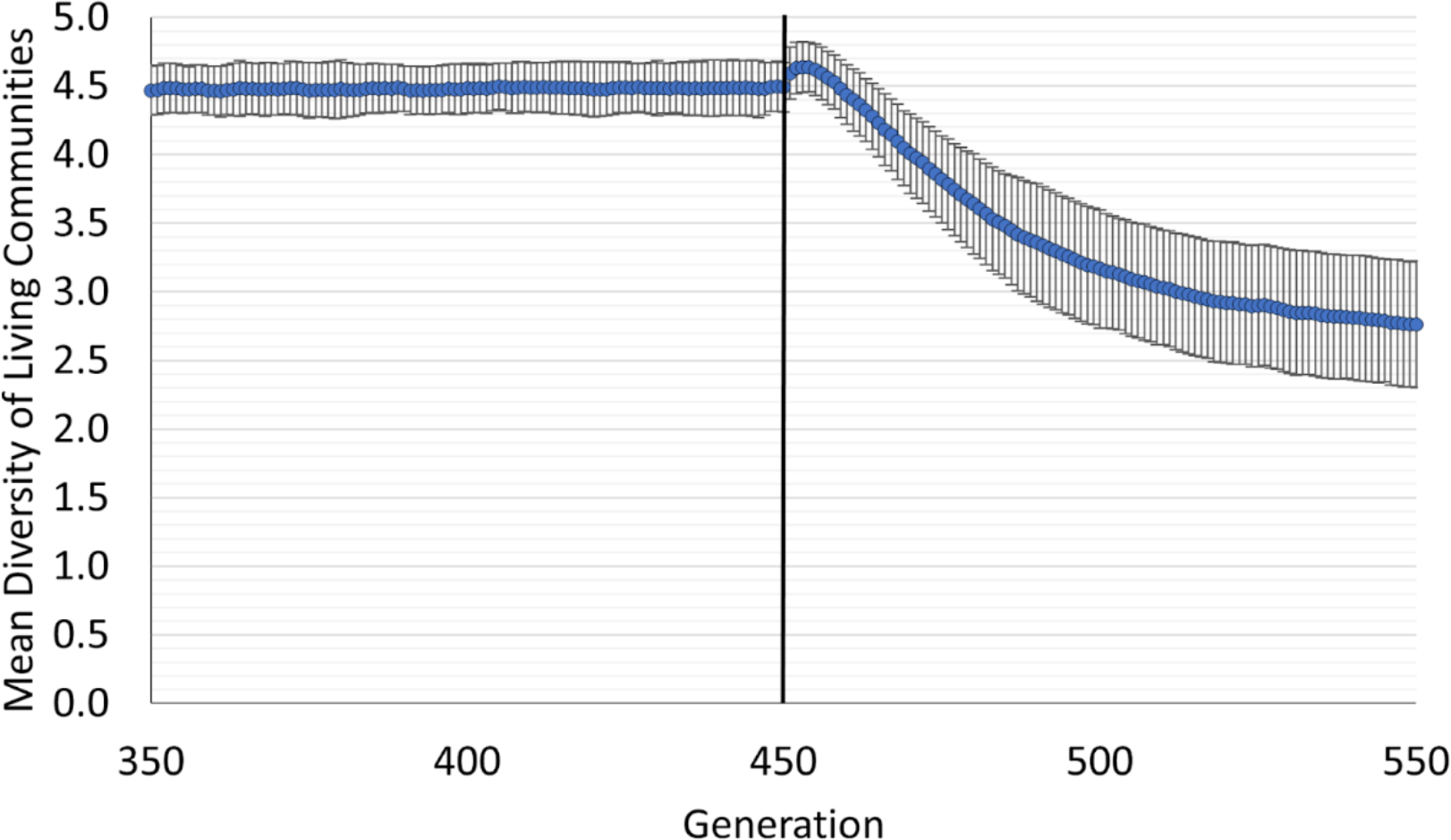
Shannon-Wiener diversity in living communities 100 generations before and after environmental change, across all 100 simulations. The initial increase after generation 450 reflects persistence of the previous high ecological quality community (i.e., priority effects) coupled with the immigration of new, low quality species. Overall changes in community composition lagged behind the rapid change in environmental conditions. The black horizontal line corresponds to the timing of environmental change. Error bars represent one standard deviation.

**Figure S9.**
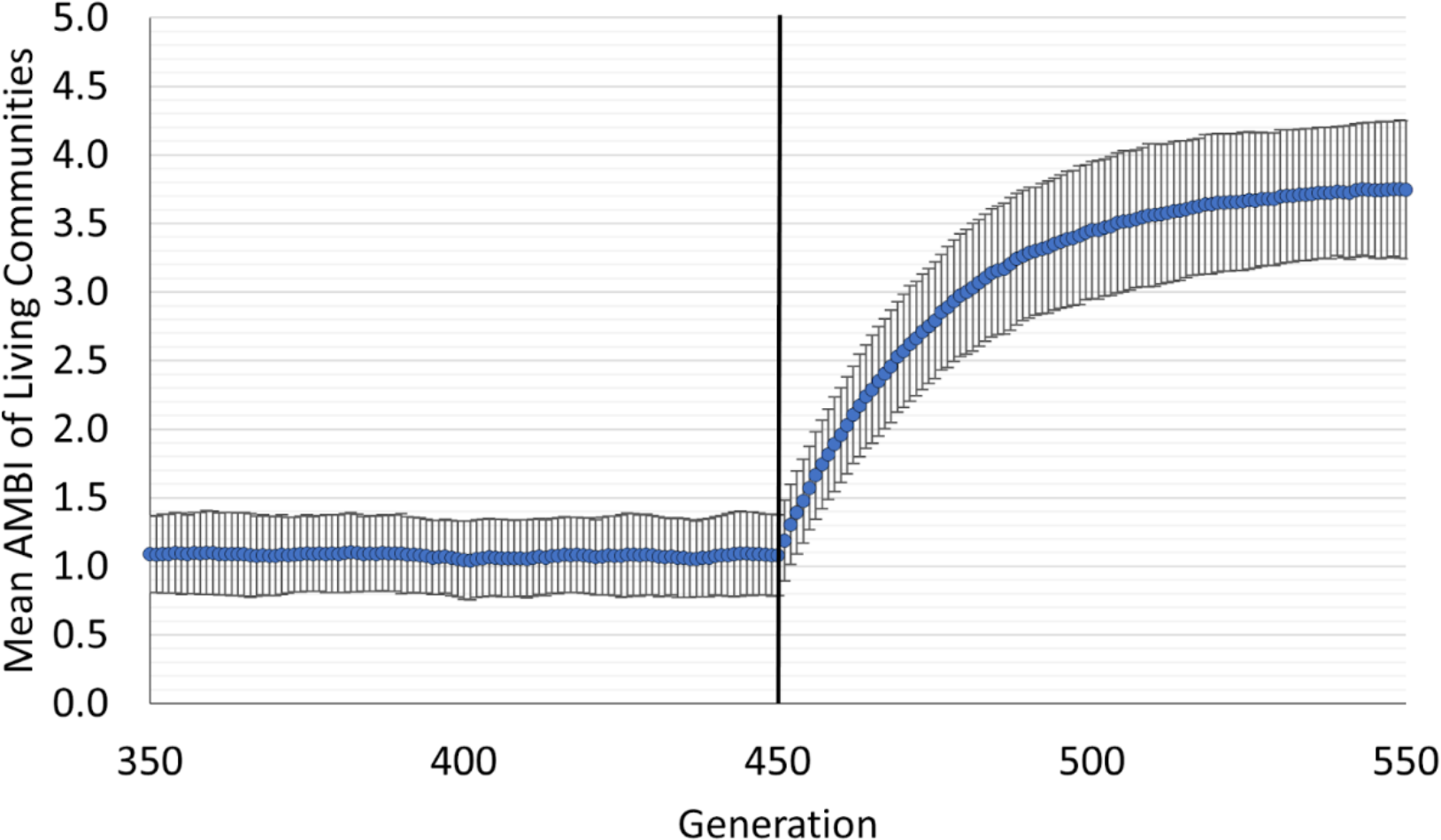
AMBI values for living communities 100 generations before and after environmental change, across all 100 simulations. Overall changes in community composition lagged behind the rapid change in environmental conditions, as high ecological quality species were gradually replaced by low ecological quality species. The black horizontal line corresponds to the timing of environmental change. Error bars represent one standard deviation.

